# The genetics of the human face: identification of large effect single gene variants

**DOI:** 10.1101/165266

**Authors:** Daniel J M Crouch, Bruce Winney, Willem P Koppen, William J Christmas, Katarzyna Hutnik, Tammy Day, Devendra Meena, Abdelhamid Boumertit, Pirro Hysi, Ayrun Nessa, Tim D Spector, Josef Kittler, Walter F Bodmer

**Author notes:** These authors contributed equally to this work (Daniel J M Crouch, Bruce Winney). Deceased (William J Christmas). **Corresponding author:** Walter Bodmer, Cancer and Immunogenetics Laboratory, Weatherall Institute of Molecular Medicine and Department of Oncology, University of Oxford.

## Abstract

In order to discover specific variants with relatively large effects on the human face we have devised an approach to identifying facial features with high heritability. This is based on using twin data to estimate the additive genetic value of each point on a face, as provided by a 3D camera system. In addition, we have used the ethnic difference between East Asian and European faces as a further source of face genetic variation. We use principal components analysis to provide a fine definition of the surface features of human faces around the eyes and of the profile, and chose upper and lower 10% extremes of the most heritable PCs for looking for genetic associations. Using this strategy for the analysis of 3D images of 1832 unique volunteers from the well characterised People of the British Isles study [1, 2] and 1567 unique twin images from the TwinsUK cohort (www.twinsuk.ac.uk), together with genetic data for 500,000 SNPs, we have identified three specific genetic variants with notable effects on facial profiles and eyes.

**Significance statement:** The human face is extraordinarily variable and the extreme similarity of the faces of identical twins indicates that most of this variability is genetically determined. This level of genetic variability has probably arisen through natural selection, for example, for recognition of membership of a group or as a consequence of differential mate selection with respect to facial features. We have devised an approach to identifying specific genetic effects on particular facial features. This should enable the understanding, eventually at the molecular level, of the nature of this extraordinary genetic variability, which is such an important feature of our everyday human interactions.

**Author Contributions:** WFB conceived the project. BW and TD organised collection of PoBI data and KH, DJMC, DM, AB and WFB assisted in data collection. TDS, PH and AN collected TwinsUK data. WPK and WJC conducted image registration analysis under supervision of JK. DJMC analysed registered image data and genetic data under supervision of WFB. WFB and DJMC wrote the manuscript, with WJC and WPK contributing additional technical material. WFB and BW supervised the project.

## Introduction

Face to face, that is how we mostly recognise each other, and we can do that because the human face is so hugely variable. Francis Galton pioneered the idea that twins could be used to study the different effects of ‘nature and nurture’ (a term he created) on various human traits [3]. In particular, having noticed that twins fell into different categories one of which was determined by their extreme similarity, which we now recognise as identical or MZ twins, he argued that this suggested that the resemblance was due much more to nature than nurture. This foreshadowed our current recognition of that the fact that, because the faces of genetically identical monozygous (MZ) twins are extraordinarily difficult to distinguish, especially at first sight, the varying facial features by which we recognise people are almost totally genetically determined. The added fact that the facial features of identical twins raised apart are as similar to each other as those raised together [4] strongly supports the view that normal environmental effects on facial features as we recognise them are usually very limited.

There is strong anecdotal evidence that similar facial features often tend to occur in families and follow on from one parent or recent ancestor to the next generation. This suggests the existence of single gene variants for such facial features, with relatively large effects. There are evolutionary arguments to support this expectation. The extent of genetic variation that must exist to explain the high level of variation for facial features has most probably been driven by natural selection, possibly connected with individual recognition and recognition of group membership. In addition, there are special regions of the brain for face recognition in humans [5] as well as in other primates [6]. The evolutionary arguments suggest that our minds are ingrained to perceive those features of a face that are likely to be strongly genetically determined. This emphasises the need for genetic studies to identify facial features with relatively high heritabilities. However, recently published studies on normal facial variation, using population based genome wide association studies (GWAS), have only found variants with quite small effects on facial features that have mostly only moderate heritabilities [7–11].

Our approach to finding large effect variants uses facial images from MZ and dizygotic (DZ) twins and a novel technique for predicting the ‘additive genetic value’ of each facial measurement, which collectively effectively determine the heritable components of the facial phenotype. The additive genetic value is a concept devised by animal breeders (see e.g. [12]), that is defined to be the expected measurement for each individual, conditioned on their genetic information. In other words, the additive genetic value is what we expect an individual’s phenotype to be if the environmental variance and measurement error were eliminated. Using additive genetic values should then give rise to measurements of facial features that are more heritable than those based on the original ‘raw’ measurements. The prediction of additive genetic values uses the genetic variances and covariances between measurements that are obtained from MZ and DZ twins with no reference to specific genotypes. Using these additive genetic values, we then base our analysis on genetically comparing individuals who lie in the extremes of the phenotypic distributions with those who do not. We used a portable version of the 3dMD camera system to obtain 3 dimensional (3D) images of 1832 unique volunteers from the very well characterised People of the British Isles study [1, 2] and 1567 unique twin images from the TwinsUK cohort, previously used to analyse the heritability of facial phenotypes derived in various ways from the camera measurements [13] (www.twinsuk.ac.uk). For most of these individuals we had genetic data for at least 500,000 SNPs. In addition, we had 33 images of East Asian (mainly Chinese) volunteers for comparison with the UK images. This enabled us to look for features in the UK faces that overlapped those in East Asians, which presumably must have a genetic basis.

Each image models the surface of the face with about 30,000 points. Using an approach that harks back to Galton’s method for obtaining average faces [14], fourteen landmark points, such as the tip of the nose, were marked on each facial image, which, through a process of mesh registration, enabled the overlaying of all images in relation to each other. This then made it possible to estimate the additive genetic values of each position on a face for subsets of the data within a facial region around the eyes and for a profile. The two resulting sets of values were then subjected to a Principle Components Analysis (PCA). We then searched for those Principle Components (PCs) that had high heritability or were notably associated with the overlap of UK and East Asian faces. At this stage, the top and bottom 10% of PoBI individuals for each of the chosen PCs were used for a genome wide association analysis. Discovery SNPs based on the UK PoBI data were then verified using the TwinsUK data. This procedure has so far yielded three convincing associations for SNPs with minor allele frequencies (MAF) around 10% in the UK population and with Odds Ratios (ORs) greater than 7 based on recessive models for the minor frequency alleles.

## Results

### 3dMD images, landmarking and face registration

The primary data for our analysis of facial features are 3D images of 1832 volunteers from the PoBI project, 1567 individuals from the TwinsUK cohort and 33 East Asian volunteers, taken using a 3dMD camera (www.3dmd.com). The images produced by the 3dMD proprietary software define the surface of each face in terms of 29,658 points in 3D space, giving 29,658x3=88,974 variables. In order to do meaningful calculations on this collection of facial images it is necessary to align the faces with each other so that any given position on a face can be matched with the corresponding position on any other face. This is done by a combination of manually annotating each face with 14 well defined landmarks (for example the tip of nose or corners of eyes) and then using a series of algorithms that register each face against a generic model face (see materials and methods and the SI for the details of these processes).

### Additive genetic value prediction and choice of facial sub-regions

As already mentioned, in order to help identify facial features with high heritability we have devised a method for obtaining the ‘additive genetic values’ (AGVs) of every point on a face. These are known as ‘breeding values’ in the quantitative genetics literature. We do this by using the MZ and DZ twin data to estimate the additive genetic variances of each point, and the covariances between points, which are needed to estimate a set of coefficients *θ* _*jk*_ using a least squares minimisation. The predicted AGV ý_*ij*_or each individual *i* at variable *j* is then given by the equation

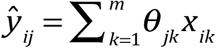

where x_*ik*_ are the observed values of the coordinates of the *k*th facial variable (an x, y or z axis position for a particular point) for individual *i* and the summation is over the total, *m*, of observations per face. (see materials and methods and SI for further details of the procedures for estimating the AGVs). Ideally this process should be carried out using all the 88,974 variables that are used to define the surface of each face. Computational limitations have so far limited us to carrying out this AGV estimation for two sub-regions of the face (eyes and profile) chosen by visual inspection. (see Fig. 1 and Materials and Methods for further details)

**Fig. 1.**
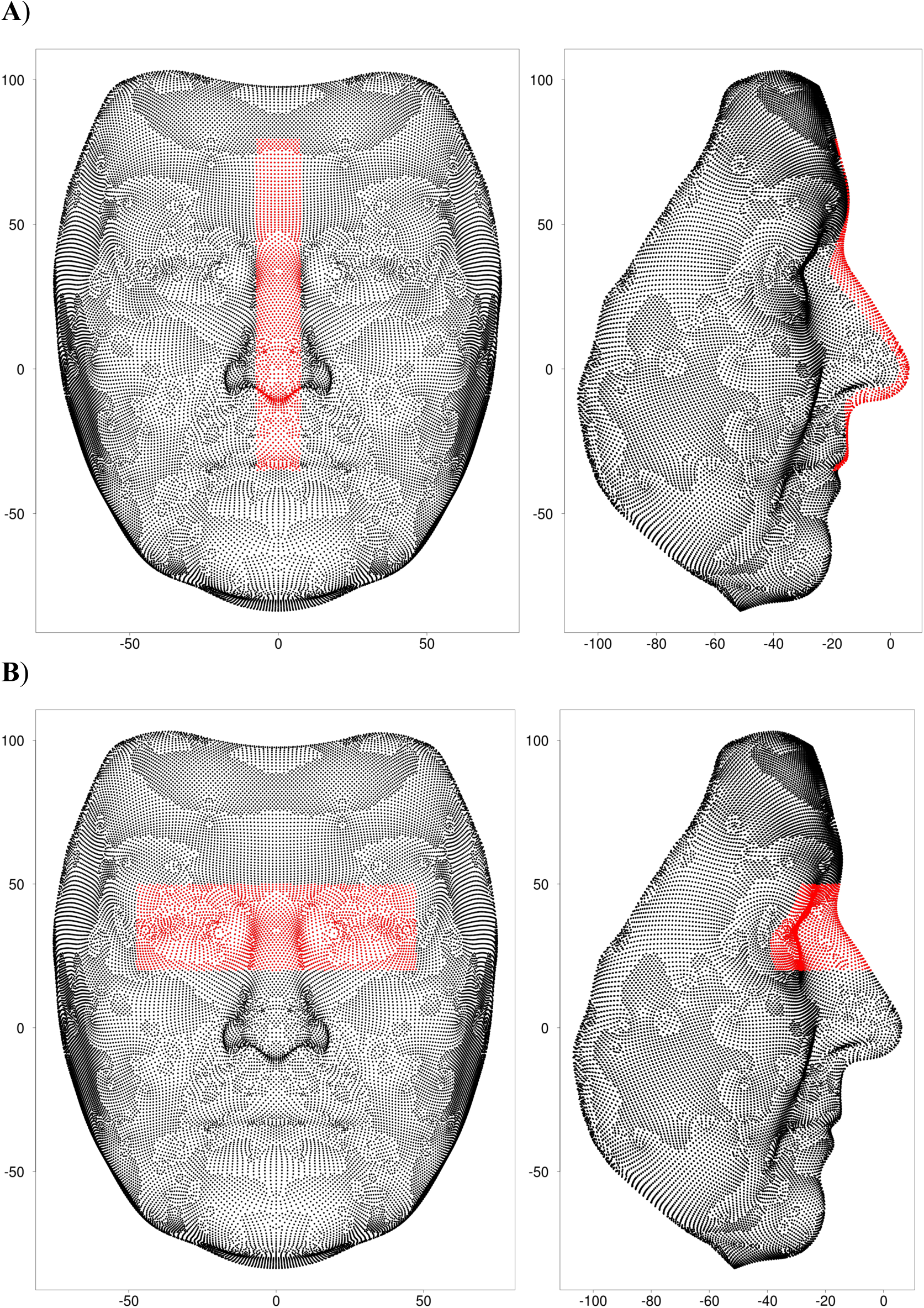
Points, highlighted in red, constituting the A) profile and B) eyes sub-regions, with the scale for both axes in mm.

The eyes sub-region includes 2736 points, giving 3 × 2763 = 8289 variables for further analysis. Only the height(y) and depth(z) dimensions were used for the profile sub-region, as the width(x) dimension was much less heritable (see Fig.S2). There are 1646 points in the profile sub-region and so 2 × 1646=3292 variables were used for further analysis.

The mean heritability of the additive genetic values for eyes was 76.1% and for the profile 81.5%, as compared to 69.8% and 76.6% respectively for the raw data. The gain in heritability for the additive genetic values for the profile is shown more directly in Fig2, which plots for all the variables, the histograms for both the raw data and the additive genetic values as determined from a subset of the TwinsUK data not used for the estimation of the additive genetic values. This clearly shows the significant increase in the additive genetic value heritabilities and also, perhaps more importantly, the substantial reduction in the variance of the additive genetic value heritabilities. This is due to much improved additive genetic value heritabilities for variables with previously very low heritabilities. (see also Fig. S3).

### Shape translation and Principal Components Analysis

As described in Material and Methods, PCAs were carried out on the additive genetic values after a Procrustes-like transformation that centered both profile and eyes sub-regions on the origin to avoid any influence of large scale craniofacial morphology causing differences between individuals in the positions of regions. The PCAs were then carried out separately for the eyes and profile facial sub-regions, using all the PoBI, TwinsUK and the 33 Asian individuals’ images combined. The resulting eigenvectors were used as the PC scores.

Five PC axes were chosen for genetic association analysis from the largest 50 PC axes, for each of the eyes and profile sub-regions. The choices were based on PC scores having a heritability of greater than 75% as determined from the TwinsUK data or having a heritability greater than 65% and a relatively large difference between the mean PoBI and East Asian scores. (see Fig. S4). This latter criterion is based on the fact that major ethnic groups generally have certain distinct types of facial features, which must be largely genetically determined. This implies that, for example, there are facial features common in East Asian populations which are much less common in European populations, and that this difference is due to one or more face-controlling genetic variants that are significantly more frequent in East Asian than European populations.

### Discovery genetic association analysis

Quantitative trait analysis generally assumes all measurements for a given trait are from the same normal distribution. If, however, there are relatively small subsets of individuals with different distributions at the upper or lower end of trait values, an overall quantitative analysis may well miss this element of data heterogeneity. Thus, in order to investigate the hypothesis that there are genetic variants conferring large effects on facial morphology, we focus on individuals with facial features that can, in some sense, be considered ‘extreme’ relative to the general population. We therefore dichotomised the PC scores for the facial phenotypes into subsets of upper and lower extremes, chosen to be the top and bottom 10% of individuals when ranked according to their PC scores. This meant that the five PCs chosen for further analysis from each of the eyes and profile sub-regions gave rise to a total of 20 extreme phenotypes to be tested for genetic associations.

Only the PoBI data were used for the SNP association discovery analysis, leaving the TwinsUK data for subsequent verification of those SNPs showing initial significance in the PoBI analysis. As described in Materials and Methods and the SI, after QC procedures there were 3161 PoBI individuals (1532 males and 1629 females) genotyped for 524,576 SNPs, of which 512,181 were non-sex linked, and were used for the association analysis. Of the 3161 PoBI individuals, only 1423 (652 males and 771 females) had both good quality facial images and genotypes, and these were used for the association analysis. There were, however, a total of 1832 imaged individuals and all of these were used for the definition of the phenotypes, as described above.

For each PC, the individuals in the 10% extreme (upper or lower) were tested against all those not in the extreme together with un-phenotyped individuals. The latter will, of course, include some individuals with the extreme phenotype being tested, which will dilute any suggested association but cannot bias the result towards an association. We assume that the relatively large number of PoBI controls who were not phenotyped offsets any slight loss of power for detecting an association due to the presence of a relatively small number of people with the extreme phenotype in the controls.

In order to allow for male versus female differences all association analyses were performed two times, namely for female extremes (excluding male phenotyped individuals - approximately 77 extremes, 2478 controls) and combined male and female extremes (approximately 142 extremes, 3019 controls). Male extremes were not studied at this stage because the TwinsUK data to be used for verification hardly include any males. In total, there were therefore 4 association analyses performed for each of the 10 PCs under consideration (females and combined sexes, each in upper and lower extremes) for the eyes and profile face sub-regions or effectively 40 different facial phenotypes.

It is important to emphasise that none of the steps taken to reach this definition of phenotypes involved any relationship to the SNP genotyping. Thus, each of these phenotypes was initially tested independently for genome wide associations, using only the number of SNPs used for the association analysis to take into account multiple comparisons for assessing significance, just as is done for standard GWAS for diseases.

For each of the selected 40 PC extremes, assessment of the significance of SNP associations was based on analysis of the 3x2 contingency tables of genotypes (*aa/Aa/AA*, where *a* represents the minor frequency allele) versus extreme/control status, for all 512,181 non-sex linked SNPs. Dominant(*A*) and recessive(*aa*) models were tested for each SNP and the P-value for the best fit then used, multiplied by 2 for the extra comparison being made. The relatively small number of extreme individuals motivated a careful selection of the appropriate statistical test for contingency table significance. This is especially pertinent when examining models of inheritance involving the effects of minor allele homozygotes, which can be at quite low frequencies amongst the extremes for minor allele frequencies of around 10%. In practice, we therefore only considered SNPs with minor allele frequencies greater than or equal to about 10%. In order to establish the most appropriate test, null hypothesis simulations of 3x2 tables were performed, and the type-I error rates quantified.

The best performing approximate test, showing only mild deflation of P-values within the borderline genome-wide significance range (10^−5^ to 10^−6^), was an implementation of a Wald test, in which standardised log odds ratios under recessive and dominant models were tested against an *N*(0,1) distribution for significant departures from 0 [15] (see Materials and Methods and SI for further details).

Variants were deemed to be significantly associated with an extreme facial feature by passing one of three thresholds:

a. Having a P value below the standard level of genomewide significance (5x10^−8^).
b. Belonging to the candidate SNP set and having a P-value below 500,000 x 5x10^−8^ / 66,769 = 3.7x 10^−7^, where 66,769 is the number of tested SNPs in the candidate list (see Materials and Methods).
c. Having a P-value below 5x10^−4^ and an odds ratio (OR) greater than 10. The rationale for this threshold is that, at the cost of a greater number of false positives, variants could be discovered with i) biologically significant consequences and ii) high power for replication, conditional on them being true positives.

By these criteria there were, for the females, 12 significant associations for the profile and 5 for the eyes. There were, in addition, for combined males and females 1 SNP association for the profile and 2 for the eyes. A total of 13 SNPs for the profile and 7 SNPs for the eyes were therefore taken forward for verification in the TwinsUK data. For the profile, 4 of the 7 phenotypes (accounting for 7 out of 13 associated SNPs) and for the eyes 5 out of 6 phenotypes (accounting for 6 out of 7 associated SNPs) with initially significant findings showed a substantial difference between East-Asian and European individuals. Interestingly, all associations were more significant under a recessive rather than dominant inheritance model.

### Verification in TwinsUK data

The TwinsUK dataset provides the basis for valid and unbiased replication since, although it was used for the additive genetic value estimation, this process makes no reference to DNA data. Of the 7 eye-associated discovery SNPs, 1 (rs2039473, eyes PC3 associated in females) was not present on the TwinsUK genotyping platform and so could not easily be pursued further. One SNP of the 13 selected for verification for the profile had an r^2^ > 0.1 with another discovery hit. Thus, retaining the SNP with the highest OR in the discovery analysis, there remained 12 SNPs with profile associated phenotypes and 6 eyes-associated SNPs to be taken forward for verification.

For each of these 18 verification SNPs, association was only tested for the PC extreme with which it was associated in the discovery analysis. To increase power, control samples from the appropriate discovery analyses were incorporated into the replication contingency tables before testing. A Wald test (1-tailed) was then performed under the appropriate inheritance model for each of the 18 verification SNPs.

A complication in using the TwinsUK cohort as a replication dataset is the high degree of relatedness between MZ and DZ twins. Removing all related individuals satisfies the requirements for independent samples required by standard statistical analyses, but also loses information and results in an arbitrarily chosen set of unrelated samples. To address these issues, we carried out a randomisation and permutation testing analysis using a limited number of the DZ twins. Thus, noting that statistical power might be gained by utilising more than half of the DZ twins per analysis, and that a pair of DZ twins contains the genetic information equivalent to 1.5 unrelated samples, we performed 10,000 random selections of one from each pair of DZ twins plus, for a random half of those DZ twins, their twin relative.

Each random selection thus included 75% of the available 327 DZ twin pairs, plus 291 composite MZ individuals and 84 remaining individuals with no relative available, which is roughly equivalent to the full set of independent genetic information available from the TwinsUK data. The appropriate genotype distributions for the extremes and non-extremes from this data set were then produced using the mean cell counts over the random selections, rounded to the nearest integer. For the controls, the PoBI controls (non-extreme phenotyped PoBI individuals and all genotyped but not phenotyped PoBI individuals) were combined with the non-extreme twins’ data. A random permutation test was then used to assess the significance of the difference between the extremes and the controls, and the OR calculated from the appropriate recessive 2 × 2 table (see Materials and Methods for further details of these procedures).

Of the 18 discovery SNPs taken forward for replication analysis in this way in the TwinsUK data set, 3 replicated with 5% or lower FDR and with ORs greater than 5: rs2045145 (in *PCDH15*), profile PC2 associated in females; rs11642644 (in *MBTPS1)*, profile PC7 associated in females and rs7560738 (in *TMEM163),* eyes PC1 associated in the combined sexes. None of these SNPs were in any of our candidate gene regions.

### Detailed analysis of the three replicated SNP associations

For each of these SNPs we show the 3 × 2 tables for the discovery analysis, the replication analysis and for the combined TwinsUK and PoBI data. The combined data tables were formed by adding the extremes results for the PoBI discovery and the TwinsUK replication, and testing the combination for significance against the same controls used in the replication analysis (namely the combined controls from the PoBI and TwinsUK sets as described above) using a Wald test. The overall OR was estimated from the 2 × 2 table for a recessive model, as is appropriate for each of the three replicated associations, and the P values were for one sided tests, given the expected direction of a SNP effect in the verification.

#### rs2045145 (PCDH15) association with the more European E x treme profile PC2 in females

The Volcano plot of – log_10_ P values against log_2_ ORs for the profile PC2 phenotype in females for the PoBI discovery analysis is shown in Fig.3A, indicating in green the rs2045145 SNP that replicated. The green lines are drawn to show the 5x10^−4^ P value (horizontal) and OR10 (vertical) thresholds above and beyond which SNPs were taken forward for replication.

(B) Distribution of PC values for the PC2 profile in UK (PoBI) and East-Asian individuals, with the East-Asian distribution magnified in size by 10 for visualisation purposes. Dotted lines show the thresholds for the upper and lower 10% of individuals.

The distribution of the PC values for the PC2 profile phenotype in UK and East Asian individuals is shown in Fig.3B. This clearly demonstrates the difference between the UK and East Asian individuals for this PC2 phenotype, which is expected based on the choice of this PC for analysis including an overall difference for this PC between the UK and East Asian individuals (see Fig.S4B and Materials and Methods). In this case, it is the lower extreme, being the more characteristically UK face, that shows the genetic association. The 3x2 tables for the rs2045145 SNP for the discovery analysis, the replication analysis and the combined TwinsUK and PoBI (i.e. replication and discovery) data are shown in Table 1.

**Table 1.** rs2045145: Association with female profile PC2 European-like extreme. 3 × **2 tables for PoBI discovery, TwinsUK verification and the combined data. P values and ORs are for minor allele (*aa*) recessive model**.

| Discovery | *aa* | *Aa* | *AA* | P-value | OR |
| --- | --- | --- | --- | --- | --- |
| Extremes | 6 | 7 | 63 | 1.01x10<sup>-4</sup> | 10.03 |
| Controls | 21 | 452 | 2004 |  |  |
| Replication |  |  |  |  |  |
| Extremes | 4 | 9 | 70 | 2.25x10<sup>-3</sup> | 6.18 |
| Controls | 25 | 615 | 2601 |  |  |
| Combined |  |  |  |  |  |
| Extremes | 10 | 16 | 133 | 2.46x 10<sup>-7</sup> | 8.64 |
| Controls | 25 | 615 | 2601 |  |  |
| Minor allele(*a*) frequency | 0.103 |  |  |  |  |

The gene frequency of the *a* allele in a representative Chinese population (301 Chinese from the 1000 genomes project [16] was 0.06, and so, as expected based on the direction of the phenotypic effect, lower than in the UK population.

For visualisation of the difference between the extreme facial phenotypes, average faces within each extreme (upper and lower 10%) were produced by plotting the arithmetic mean for each coordinate measurement in each vertex, among all individuals falling into the designated extreme, and overlaying with a surface texture. Only females were used to produce average faces, so as not to obscure the genetic difference by sex differences. Average faces were produced using all Asian females and all PoBI females separately. The average profiles for the East Asian females and the upper and lower 10% of the PoBI females for the PC2 profile are given in Fig.4. It is the lower more European looking extreme profile of Fig.4C that is significantly associated with the recessive (*aa)* genotype of the rs2045145 SNP with an overall OR of 8.64. It is, however, notable that the East Asian alike PoBI female face (Fig.4B) has an upturned nose and upper lip, and a receded chin, similar to the average East Asian female face.

The rs2045145 SNP is in an intron of two trancripts (*PCDH15-220* and *PCDH15-225*), 670Kb downstream from the fist exon of *PCDH15-225* and 580Kb upstream from its terminus and so itself has no obvious functional effect. Using the approach to functional analysis given in Materials and Methods, three potentially functionally relevant variants, but with relatively low ORs, were found to be in LD with rs2045145 (r2=0.53, 0.53 and 0.55). These were within 5Kb upstream of a lncRNA (207bp mature transcript) situated in an intron of *PCDH15*. *PCDH15* is one of several genes for which loss of function recessive variants cause one of the forms of Usher syndrome, which involves hearing and balance malfunction in addition to retinal defects. Usher syndrome is not usually associated with a distinctive facial appearance, although dysmorphology has sometimes been seen [17]. *PCDH15* is expressed in the nose (olfactory epithelium and nasal cartilage) of developing mice [18], perhaps fitting in with the observation that the minor homozygote is associated with a straight rather than upturned nose (Fig. 4C). An additional tagged SNP with an OR of 4.65 in the sequenced TwinsUK data (rs61850893, r^2^=0.72) is situated 229bp upstream of a conserved intron region within *PCDH15*, but it’s possible functional significance is not clear. As shown in FigS9A, the discovery SNP lies in a region of sequence conserved between primates at a position where the common allele (*G*) is the common allele in humans. This suggests that in this case, the discovery allele could itself be functional with respect to the facial phenotype.

**Fig. 4.**
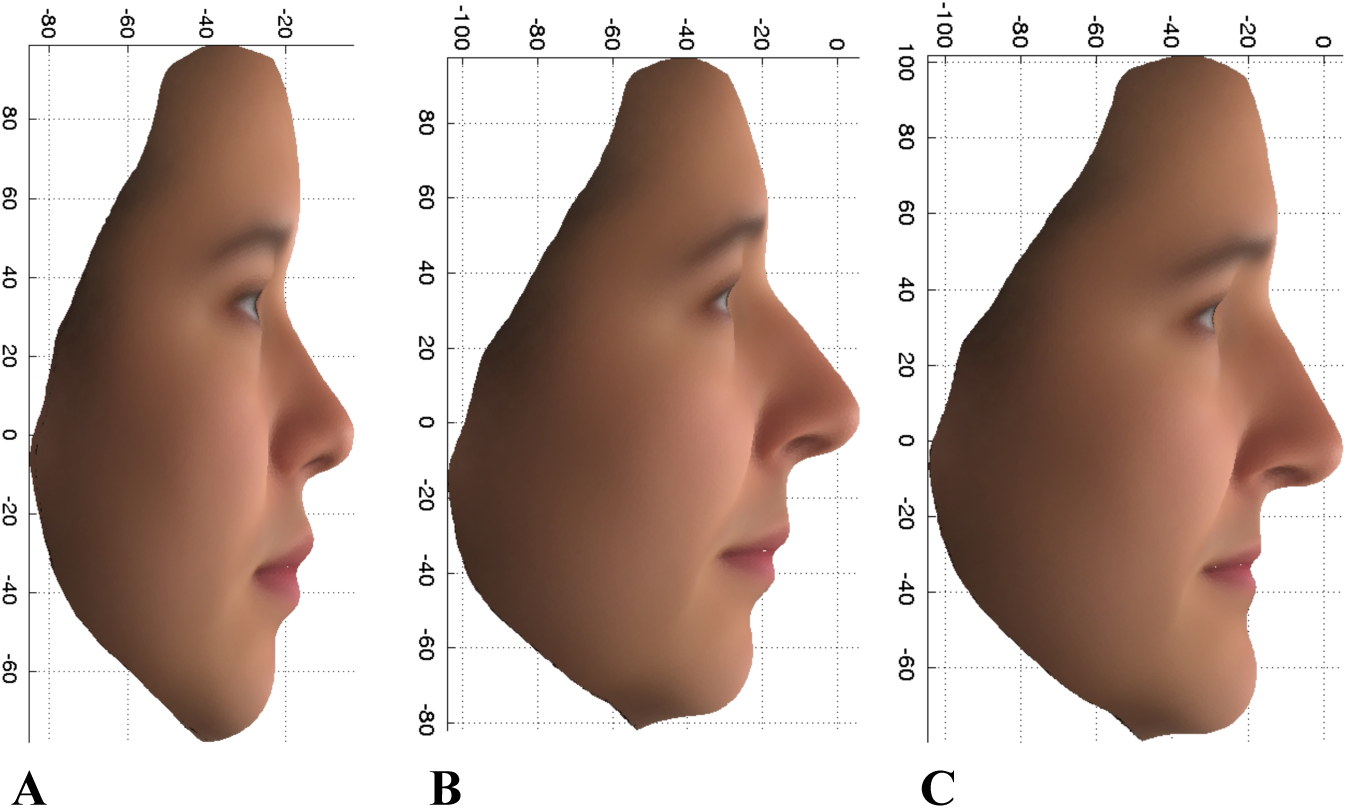
PC2 profile: Average faces for 14 East Asian females (A), and the upper 10% (more East-Asian) (B), and lower 10% (more European) (C) extremes of the PoBI females.

#### rs11642644 (MBTPS1) association with upper extreme profile PC7 in females

Table 2 shows the 3x2 associations for rs11642644 for the discovery, replication, and combined data analyses and Fig. S6A, the volcano plot that led to the choice of this SNP for replication.

**Table 2.** rs11642644 (*MBTPS1*) associations with profile PC7 upper extreme in females.

| Discovery | *aa* | *Aa* | *AA* | P-value | OR |
| --- | --- | --- | --- | --- | --- |
| Extremes | 6 | 15 | 56 | 3.20x10<sup>-5</sup> | 11.55 |
| Controls | 18 | 459 | 2001 |  |  |
| Replication |  |  |  |  |  |
| Extremes | 3 | 13 | 69 | 1.10x10<sup>-3</sup> | 5.58 |
| Controls | 22 | 569 | 2632 |  |  |
| Combined |  |  |  |  |  |
| Extremes | 9 | 28 | 125 | 1.31x10<sup>-6</sup> | 8.56 |
| Controls | 22 | 569 | 2632 |  |  |

Minor allele(*a*)frequency 0.097
P values and OR are for the minor allele (*aa*) recessive homozygote

**Fig. 5.**
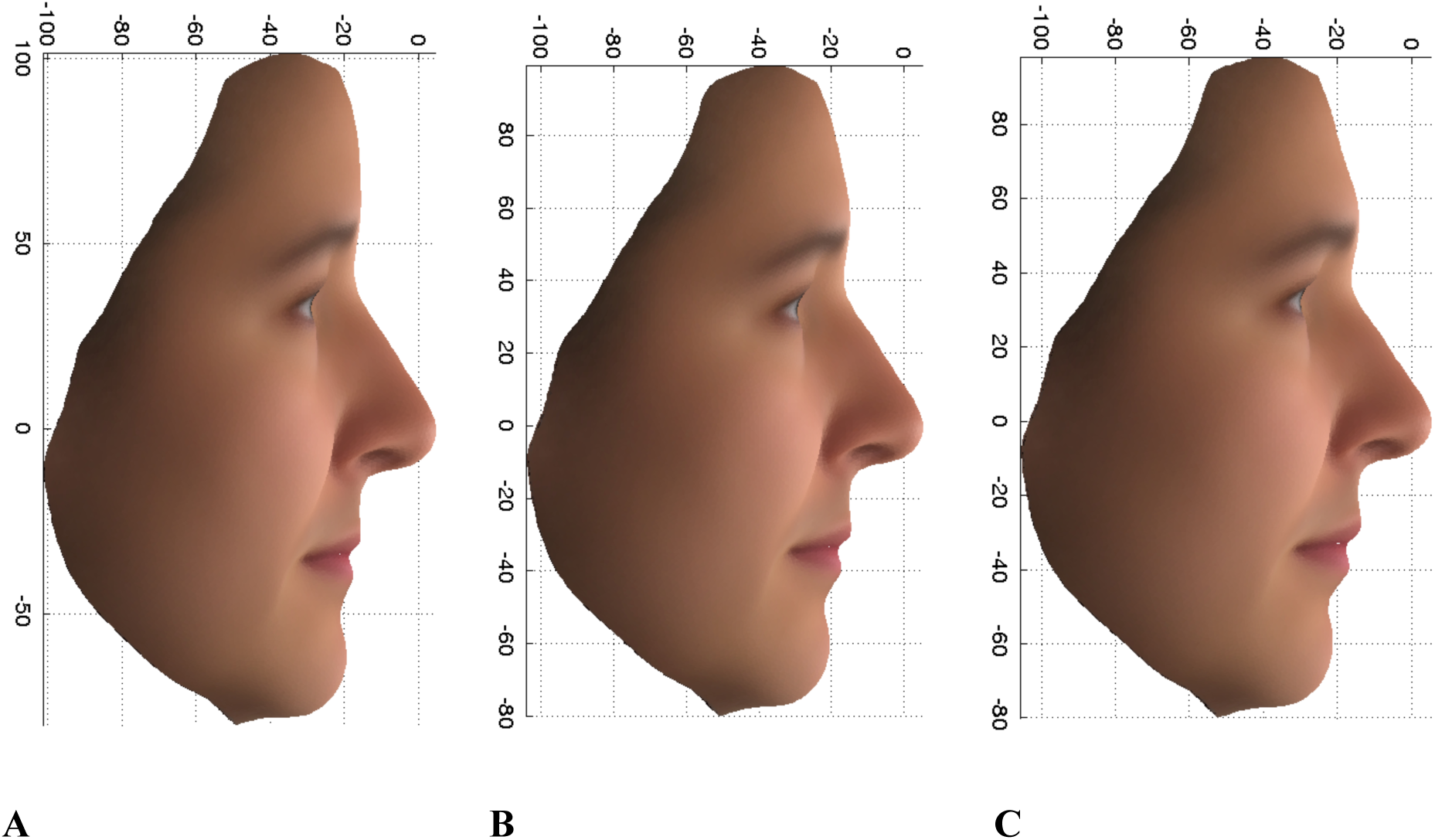
Average profile PC7 female faces for the upper (A) and lower (C) extremes and the overall average (B).

In this case, it is the upper extreme, Fig. 5(A) that is associated with the recessive homozygote(*aa*) for the rs11642644 SNP, and this does not show any obvious association with the East Asian faces (see Fig. S6B), as expected from the data in Fig. S4B on which the choice of profile PC7 was made for further analysis.

The SNP rs11642644 is in the gene *MBTPS1* and is located in an exon of a 543bp transcript (*MBTPS1-016*) which is within the *MBTPS1* gene. This exon has no open reading frame and is likely to be part of a lncRNA. Two of the transcript’s three exons overlap with those of the protein coding *MBTPS1* transcript. rs11642644 also lies in a region of open chromatin. The minor allele (C), associated with the upper 10% extreme phenotype as a recessive genotype, is present in African Green Monkey, Macaque and Olive Baboon, while the major allele (T) is present in Orangutan, Gorilla, Chimpanzee and Marmoset (Fig S9B). This suggests that the minor allele is itself a functional candidate, as these species differ considerably in their craniofacial morphology.

A major function of *MBTPS1*, also known as site-1 protease (*SP1*), is to cleave *SREP* proteins in the endoplasmic reticulum [19]. The *SREP* proteins are transcription factors encoded by two genes, *SREBF1* and *SREBF2*, which regulate production of enzymes responsible for steroid biosynthesis. *SREBF1* is in a 3.7-Mb region of 17p11.2 which is reciprocally deleted and duplicated in Smith-Magenis and Potocki-Lupski Syndromes [20], both of which have accompanying facial dysmorphias. *SREBF2* is within the 22q13 deletions in Phelan-McDermid syndrome (also known as 22q13 deletion syndrome), also with a dysmorphic facial appearance. Both proteins are strongly craniofacially expressed in mice at 10.5days post conception, and *MBTPS1* also seems to be expressed in a smaller region at the tip of the snout at this stage of development [21]. Taken together, this evidence provides a possible explanation for the functional effect of the rs11642644 SNP either through an effect on the open chromatin or through an effect of the presumed lncRNA on the expression of the *MBTPS1* gene.

#### rs7560738 (in TMEM163) association with the upper extreme PC1 eyes phenotype in the combined sexes

Table 3 shows the data on the rs7560738 association with the upper extreme PC1 eyes phenotype in the combined sexes for the discovery, replication, and overall analyses. The volcano plot that led to the selection of this SNP for further analysis is shown in FigS7A, and FigS7B shows that there is no East Asian association for this SNP, as expected from FigS4A.

**Table 3.**
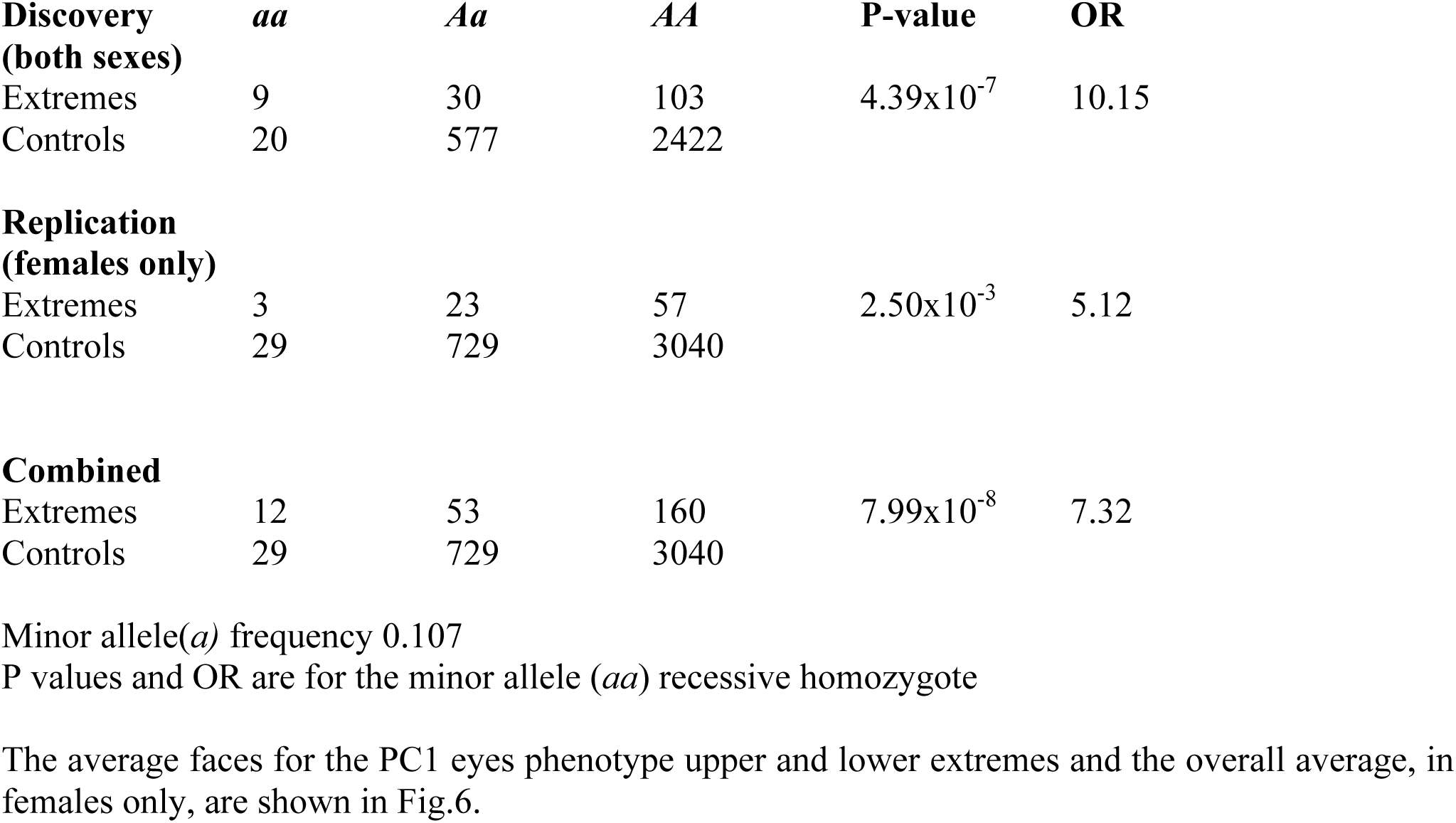
rs7560738 (in TMEM163) associations with the eyes phenotype PC1 upper extreme in the combined sexes.

It is the upper extreme (Fig6A) that is associated with the minor allele homozygote(*aa*). Note the differences in eye width and eye height, against the x and y scale respectively, which are both greater in the upper extreme. rs7560738 lies in *TMEM163*, which is a highly-conserved gene in mammals coding for a trans-membrane protein that is a putative zinc transporter expressed in the brain and retina, as well as in a limited number of other tissues. Recent work [22] shows that *TMEM163* is a binding partner of *MCOLN1*, for which loss of function mutations (including deletions and point mutations) cause Mucolipidosis type IV (MLIV) [23], a lysosomal storage disorder. Although MLIV is not commonly thought of as having distinctive facial features [24], several cases have been identified which share particular facial dysmorphias, especially around the eyelids [25, 26]. Photographs of these cases document some atypical eye shapes and positions [25]. *TMEM163* may be involved in the pathology of the disorder through its influence on cellular zinc levels, which are elevated in MLIV [27] and increased when *TMEM163* is knocked down [22]. Two orthologues of *MCOLN1* are expressed in zebra fish eyes during development, supporting the case for the *TMEM163-MCOLN1* interaction playing some role in the determination of eye morphology [28]. These data suggest that *TMEM163* could be a plausible functional candidate for the eyes PC1 phenotype. This is supported by the data on the DNA sequences around the discovery SNP in a range of primates (see FigS9C), which shows that the discovery SNP is a variant in the midst of a conserved region. It is the derived allele (A) that has an approximate frequency of 10% in Europeans

## Discussion

Our success in finding specific single SNPs with relatively large effects on facial features depends on our ability to identify facial phenotypes that have the high heritability expected from the extraordinary facial similarity between monozygous twins. Key to this has been the estimation of the additive genetic values of points on the surface of the face using data from the 3dMD camera system’s three dimensional facial scans of MZ and DZ twins. Another contribution to the phenotype heritability is the use of ethnic differences, in our case between the UK and East Asian populations. These differences are necessarily largely genetic and must be due to differences between ethnic groups in the frequencies of genetic variants with defined effects on facial features. The final contribution to our success has been the use of dichotomised variables for the genetic association analysis rather than a traditional quantitative variable approach. Basing our analysis on facial variation within the relatively homogeneous UK population has necessarily to some extent limited the range of facial genetic variability available for studying. However, the use of the well characterised and carefully sampled PoBI population as our control for genetic association studies has minimised the possibility of population stratification effects that are especially troublesome for studies on mixed ethnic populations. It is important to emphasise that none of the approaches to the choice of phenotypes to study were in any way dependent on the genetic information. This prevents any bias of the chosen phenotype towards having significant associations with SNPs, so that the correction for multiple testing of each chosen phenotype will therefore only depend on the number of SNPs tested for.

The choice of extreme phenotypes for study in order to maximise the chance of finding variants with relatively large effects, inevitably limited the number of individuals available for a genetic association analysis. This problem turned out to be particularly relevant given that the genetic associations we have found involve homozygotes for relatively low frequency variants. This led to a careful consideration of the appropriate statistical test to use for selection of variants for subsequent verification. Simulations of expected distributions using random permutations between controls and cases strongly suggested that the conventional Chi square tests were too permissive for the balance of cases and controls in our study (Fig. S5B). This led us to adopt a Wald test based on standardised log odds ratios. We also decided to take into account the estimated magnitude of the effect of a potential discovery SNP, choosing a lower threshold of an odds ratio of around 10 as a criterion for taking forward a SNP for verification, while allowing for a less stringent P-value threshold. Once a criterion has been set for the choice of SNPs to be taken forward for verification, it is then only the number of these chosen SNPs that needs to be taken into account in assessing an acceptable P-value for verification.

Another limitation of our study has been the resource limited availability of a suitable sample of phenotyped and genotyped individuals for verification. We were fortunate to have the TwinsUK samples for this, but that created the problem of effectively only having females for verification. It also posed the challenge of how best to use the information from the DZ twins without losing information due to using only one member for each pair in the verification analysis.

The three replicated SNPs were identified as significant by an association with individuals in a 10% extreme of selected PC distributions. We sought for more subtle effects by looking at the frequencies of the associated homozygotes (*aa)* and heterozygotes (*Aa)* in successive 5% quantiles across the appropriate PC distributions for combined PoBI and TwinsUK data (see Materials and Methods), as shown in Figure 7. This shows that in each case the frequency of the *aa* homozygotes is largely concentrated in the extreme 5% or 10% of the PC distribution with no obvious variation in the (*Aa)* heterozygote frequencies over the different levels of the PC7 profile and PC1 eyes phenotypes. This is consistent with the lack of heterozygote association with these phenotypes in the data given in Tables 2 and 3, and so a clearly recessive control of the extreme phenotypes. For the PC2 profile there is a remarkable association between the presence of *aa* homozygotes and the absence of *Aa* heterozygotes in the lower SNP associated extreme (OR 15.4, Yates corrected Chi square 53.2). This implies that the presence of a single *A* allele in this case positively inhibits the expression of the lower extreme phenotype. This is consistent with the observed negative correlation between heterozygotes *Aa* and the lower extreme phenotype shown in Table 1 (OR 0.51, P =0.01 Fisher’s exact test.)

**Fig. 7.**
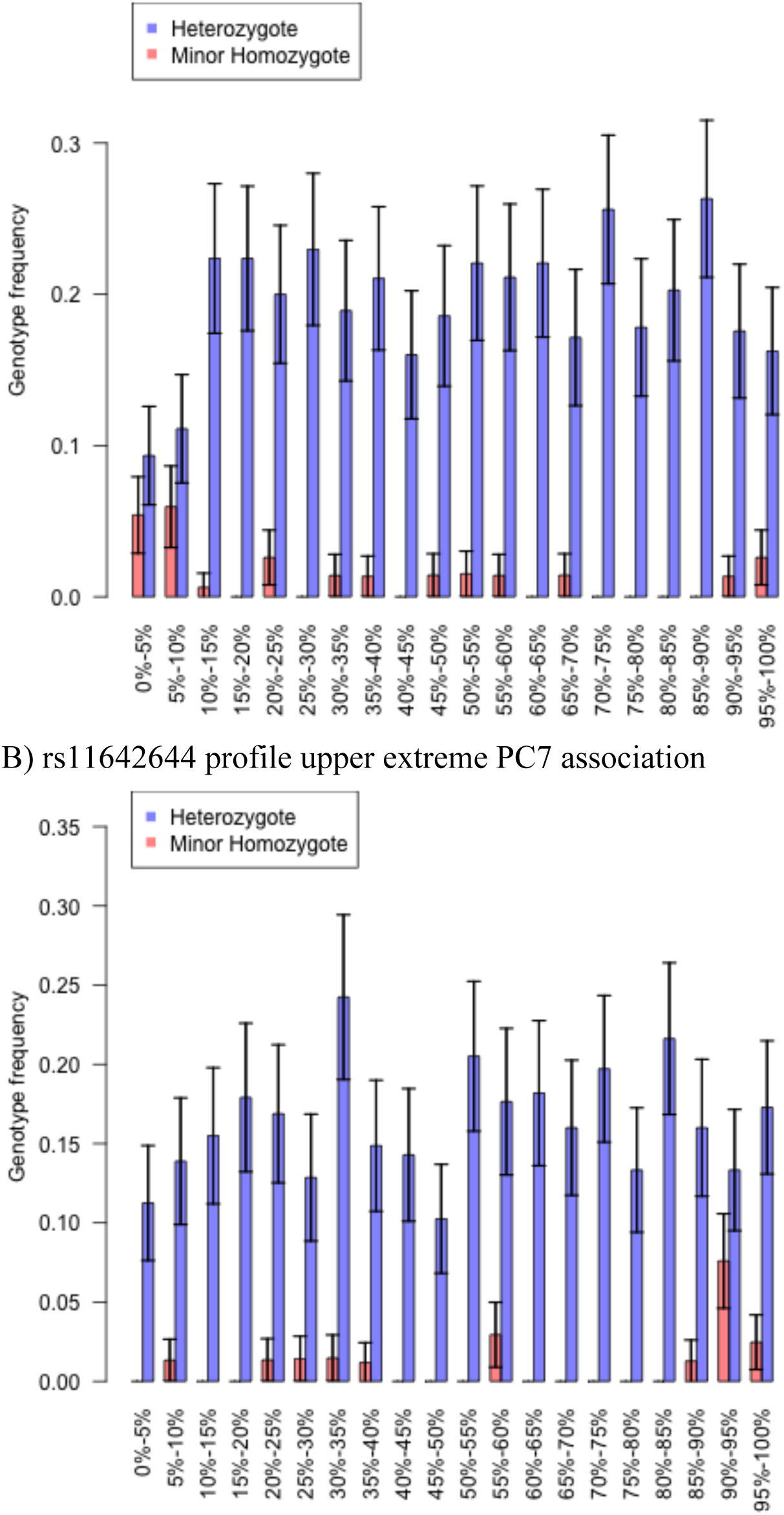
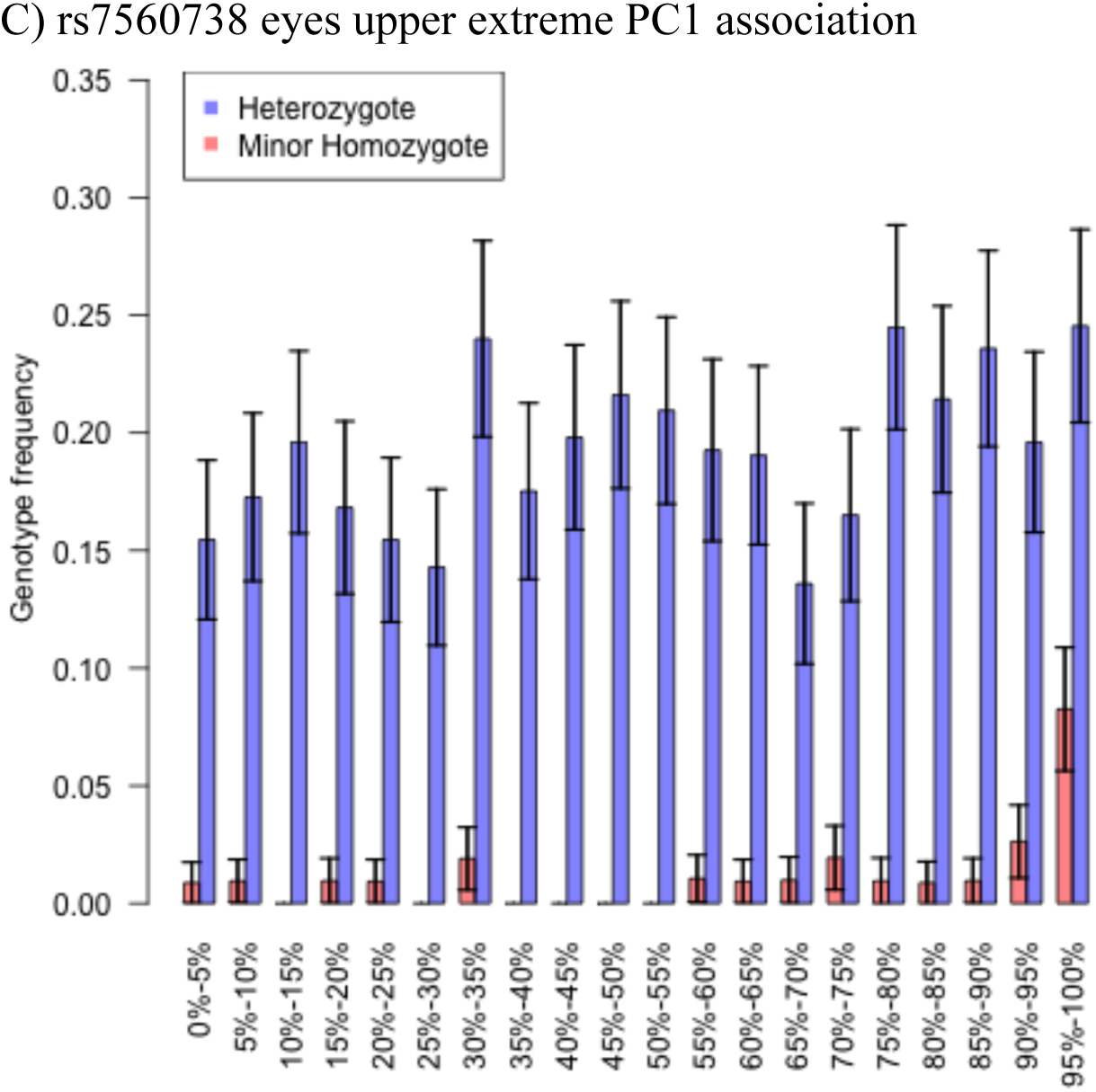
Bar graphs of frequencies of minor allele homozygotes and heterozygotes at 5% quantiles for the three verified SNP associations.

In spite of the strong recessive effects of the three verified SNPs on their associated extreme phenotypes, there are in each case many extreme phenotypes that are not recessive homozygotes for these SNPs. While some of these may be due to measurement error in the phenotype, it seems more probable, given the high heritabilities involved, that most are likely to be associated with other sources of genetic variation that our analysis was too weakly powered to detect. These could include lesser effects of either homozygotes or heterozygotes for other SNPs, or epistatic effects. The preponderance of recessive phenotypes may be due to the fact that we are looking for unusual phenotypic differences, with overall dominant effects controlling the corresponding commoner facial features.

The familial pattern of apparent parent to offspring passing on of particular facial features will be expected to be seen largely in the matings between the strongly associated homozygotes (*aa)* and the heterozygotes (*Aa*). This is because the probability of the *Aa* heterozygote*s* expressing the associated feature is 5 to 10-fold less than for the *aa* homozygotes. Since the proportion of *aa* homozygotes in these meetings will be 0.5, this gives the appearance of an apparent dominantly determined phenotype.

In our small sample of three verified SNPs there is no evidence of any particular pattern of gene functions. This is in marked contrast, for example, to the genetic control of tissue type incompatibility by the HLA system in humans or of taste and smell differences by the extraordinary complex of olfactory receptors. Selection for genetic variation in facial features connected, for example, with recognition of group membership and mate choice preferences is likely to be frequency dependent. This can explain high levels of polymorphism, as was suggested many years ago for the HLA system [29]. However, in the case of the face, the genetic variation selected for may be adventitious with respect to gene function in the sense that there may be very many different ways in which variation in particular genes can influence the inheritance of facial features.

The ultimate test for the validity of any particular SNP found to be strongly associated with a particular human facial feature will be to verify the functional effect associated with that SNP. This can be done, for example, by evolutionary comparisons relating SNP variability to facial features, such as for the differences between primate species (as we have done for our replicating SNPs) or in the variety of dog breeds, and by exploring the effects of specific gene mouse knock outs or knock ins on snout morphology.

Our studies as described here have been limited by:

i. The need for observations on larger panels of individuals, both for discovery and verification, to increase the power for detection of variants with lower gene frequencies than 0.1 and also somewhat lower ORs;
ii. The need to extend the computational tractability of the analysis in order to process the whole face; and
iii. The possibility of studying a wider range of ethnic differences.

We suggest that many more specific and relatively large genetic variant effects on human facial features will be found in the future using approaches such as we have described.

## Materials and Methods

### Sources of volunteers

The PoBI individuals were sampled from throughout the UK primarily from rural areas and from individuals all of whose grand parents came from roughly the same area, as described by Winney et al (2). The majority of the samples (2039) have been subject to a detailed population genetic structure analysis [1], which, while revealing extraordinary genetic structure related to geography, showed that the average *F*_*ST*_ between regional groups was only 0.0007. Thus, for the purpose of genetic association studies this is an ideal population. MZ and DZ twins were from the TwinsUK cohort (http://www.twinsuk.ac.uk).

### Image phenotyping

All facial images were taken using a portable version of the static 3dMDface(tm) System3dMD camera and its accompanying dedicated software (www.3dMD.com). The system uses two stereo camera units (each containing two stereo cameras, one above the other), mounted at approximately 45^o^ to the left and right of the participant, roughly a metre away. Stereo triangulation algorithms match surface features recorded by each unit, yielding a single 3D surface. Additional cameras on each unit produce colour photographs that are merged and matched to the 3D surface by another algorithm, producing the ‘texture’ map, which, for example, portrays skin and eye colour.

Where possible, 3 images were taken of each participant, to allow averaging of measurements over multiple images, with the aim of reducing technical noise. Volunteers were asked to assume a neutral, relaxed facial expression. 2556 images of PoBI individuals and 2025 images of TwinsUK volunteers were collected. Of the 2556 images of PoBI volunteers, 1832 were of unique individuals (819 male and 1005 female and 8 for whom sex was not correctly recorded), while for the twins 1567 were of unique individuals (1551 females and 16 males). In addition, single images were taken of 33 East Asian Chinese origin volunteers (14 females and 19 males). Images from all datasets were registered at 29,658 locations on the facial surface, each referred to as a vertex or point. Every vertex has 3 coordinate positions describing its x, Y and Z dimensional positions, yielding 88,974 phenotypic variables per individual.

### Facial registration

The 3D face images generated by the 3dMD camera system are provided in the form of a triangulated mesh (Fig. S1A). Each vertex has associated with it a 3D location and an RGB (red green blue) appearance value. However, the identity of each mesh vertex or point at this stage is unknown, and the number of points varies from image to image. This is dealt with by mesh registration, which involves fitting the mesh of a generic face model to the unknown mesh to produce a standardised triangulated mesh in which the identity of each node in the mesh is known (Fig. S1B; see [30, 31]). The starting point for the process is to ‘landmark’ each face manually at 14 predefined salient landmarks (Fig S1C). Landmarking is performed by 3 separate researchers for each face using the average over 3 landmarks to minimize measurement error. Following this, a series of fitting and matching algorithms is used that progressively align the generic model with the unknown input face, based initially on the landmarks, in such a way that eventually any given position in a face can be matched with the corresponding position in any other face. It is, thus, this registration process that enables meaningful calculations on the collection of facial surfaces to be made (For further details of the process see SI).

#### Additive genetic value prediction (AGVP)

The raw facial surface measurements x_*ij*_ are continuous random variables describing a position in one-dimensional space (three measurements for each vertex), where *i* and *j* are indices, respectively, for individuals (total *n*) and variables (total *m* = 88,974). We aim to estimate the unobserved values *Y*_*j*_ = *E*_*e*_[x _*j*_ | *g*], where the expectation is taken over the stochastic environmental effects (*E*_*e*_), for a given individual, for each measurement, *j*, where x _*j*_ is the random variable of which x_*ij*_ is a realisation, and *g* represents a random vector of genotypes belonging to the individual. In quantitative genetics, the departure of *Y*_*j*_ from its population mean would be termed the *genetic value.* Under purely additive genetic effects (no dominance or epistasis), the assumption we rely on below, this is the *additive genetic value*. The objective is to predict *y*_*ij*_, the realised additive genetic value for individual *i*, at measurement *j*, using the complete set of measurements taken on the same individual: {x_*ik*_ :*k* ∈1,2,3…*m*}. This is performed for each *j* in turn. Each x_*k*_is modelled as

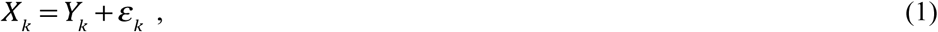

for all *k* = 1,2,3…*m*, where *ε*_*k*_represents the effects of the environment. We assume that *E*_*e*_[*ε*_*k*_ | *g*]= 0, equivalent to there being no gene-environment interactions. To estimate the coefficients θ _*jk*_, which effectively measure the predictive influence of variable x_*k*_ on *Y*_*j*_, we minimise the expected least squares error,

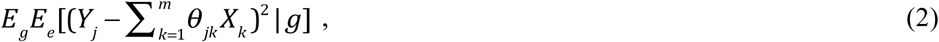

with respect to each coefficient θ _*jk*_. This double expectation is taken with respect to the stochastic environmental and genetic effects (*E*_*e*_ and *E*_*g*_respectively). After applying constraints to avoid bias and so introducing a Lagrangian parameter, and estimating genetic variances and covariances from the twins’ data assuming only additive genetic effects, we obtain a matrix equation whose solution gives the estimates of the coefficients θ _*jk*_. The predicted additive genetic values, for each individual *i* at measurement *j*, are then

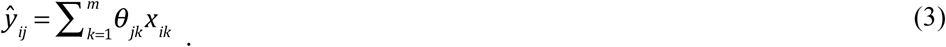

For further details of the estimation procedure see SI.

### Fitting process using facial subsets

Ideally the linear predictor uses all facial variables {x_*ik*_ :*k* ∈1,2,3…*m*} as described above, where *m* (=88,974) is the total number processed and registered from the camera system. However, computational limitations have so far prevented this, and so we used a workaround whereby analysis is restricted to two rectangular sub-regions of the face, which we call ‘profile’ and ‘eyes’. These were selected based on visual inspection of an image of the average face (Figure 1). Vertices that fall within the sub-region on the average face are taken forward for AGVP analysis in all individuals.

The eyes sub-region contains 2763 vertices, each in 3 dimensions, giving 3 x 2763 = 8289 variables for analysis, whereas the profile sub-region contains 1646 vertices. However, as the x (width) dimension of the profile is both poorly characteristic of appearance and has low heritability (Supplementary information figure S2A), only the Y (height) and Z (depth) dimensions were used for the profile, giving 2 x 1646 = 3292 variables for analysis. A process of 5-fold cross-validation was used for optimal choice of a tuning parameter in order to avoid over-fitting during AGVP analysis. (See SI). All subsequent statistical analyses were performed on the predicted additive genetic values.

### Shape translation and Principal Components Analysis

Using a Procrustes transformation approach, we translated each individual’s eyes sub-region by subtracting the mean x coordinate for their own sub-region (taken across all vertices in that region for the particular individual) from the x coordinates of all variables in the sub-region, and similarly for the y and z coordinates for both eyes and profile sub-regions. This has the effect of centering all individuals’ sub-regions on the origin. Downstream statistical analyses using these centered values are not then unduly influenced by large-scale craniofacial morphology. In order to avoid losing information that could be of genetic interest, no other Procrustes transformations, for example matching sub-regions by rotation or scaling by size, were applied. PCA was carried out in the standard way, performing eigenvalue decomposition on the *n*x*n* covariance matrix of scaled and centered facial variables for both sub-regions separately, using all TwinsUK, East Asian and PoBI images combined. The resulting eigenvectors were used as the PC scores.

Projecting TwinsUK individuals onto PoBI-fitted PC axes showed that PC scores for the largest 5 axes had r>0.9 with the scores for both sub-regions produced using the combined data. This relatively high correlation between the two sets of PC scores justifies the use of the combined PCA analysis. PC scores were averaged over all TwinsUK and PoBI individuals for whom multiple images were available, in order to reduce sources of environmental variation such as differences in facial expression and small measurement errors.

### Principle components subset selection

The largest 50 PC axes were inspected for promising phenotypes using two criteria: a) Heritability (assuming no dominance variance (see [32]) of each PC axis as determined using the PC scores of the 1567 TwinsUK individuals and b) The squared difference between the mean PoBI (European) PC score and the mean East Asian score, taken as a ratio against the within-population variance. Plots of these statistics are shown in Figure S4. PCs with heritability >0.75 or heritability>0.65 and a between/within population variance ratio >1 were taken forward for genetic association analysis.

### Genotype quality control and genotyping

Genotyping of the PoBI discovery samples was performed on two separate Illumina platforms. As part of the Wellcome Trust Case Control Consortium 2 (WTCCC2) study [33], 2,912 samples were typed on the Illumina Human 1.2M-Duo genotyping chip, 2,039 of which were previously used to analyse the population structure of the British population [1]. Since 2011, 823 newly recruited samples were genotyped on the Illumina Infinium OmniExpress-24 BeadChip (750K) platform. The intersection between the two platforms was 547,863 SNPs. Prior to QC there were 3,735 genotyped DNA samples, constituting 3,616 unique individuals.

Genotype quality control procedures were performed using PLINK versions 1.07 [34] and 1.90 [35] in combination with R scripts. Markers were removed due to showing evidence for a departure from Hardy-Weinberg Equilibrium using a cutoff of P ≤ 10^−4^. A number of samples otherwise passing QC were removed due to close relatedness or due to being duplicates of the same individual. As genotyping was performed in 7 different batches, 2 × 3 chi-square tests were performed in turn between each batch and all other combined batches, for each SNP, to detect possible batch effects. Using a P-value threshold of 10^−6^, 5024 SNPs were found significantly associated with genotyping batch, and were consequently removed. After QC, genotypes for 3161 individuals (1532 male, 1629 female) and 524,576 SNPs were retained for association analysis.

TwinsUK genotype data were available on two platforms; 1278 samples represented by 2,287,998 array-typed SNPs and 612 whole-genome sequenced samples (19,725,734 autosomal variants). Separate QC procedures were performed for the two platforms before merging. After merging the array and sequencing data and retaining variants common to both sets, there were 1,887,250 variants typed on 1794 samples, 1275 of whom were unique individuals.

For more details of quality control procedures see the SI.

### Candidate SNP list

Just under 400 candidate genes and regions were chosen by reviewing the literature on human facial dysmorphias and on facial morphologies in other species, as well as through searches of the Online Mendelian Inheritance in Man (OMIM) and Ensembl databases. SNPs were designated candidates if they were located within 75Kb of a candidate region. This yielded a list of 66,769 candidate SNPs.

### Selection of appropriate statistical test

The relatively small number of extreme individuals motivated a careful selection of the appropriate statistical test for contingency table significance. This is especially pertinent when examining models of inheritance involving the effects of minor allele homozygotes, which can be quite rare even when an allele is common. In order to establish the most appropriate test, null hypothesis simulations of 3 × 2 tables were performed, and the type-I error rates quantified. Previous work [36] has demonstrated that, when the numbers of observations are small and very low P-values are required, e.g. less than 5x10^−8^ for genomewide significance, a variety of traditional frequentist tests are conservative, either due to asymptotic assumptions breaking down, or due to inherent conservativeness in the case of Fisher’s Exact test. Simulations performed under our study design, which, in contrast with Bigedli et al (2014) had unequal proportions of extremes versus controls, it was found that most of these tests are anti-conservative. Fisher’s exact test controlled the type-I error well in simulations, but was not chosen as it is often conservative and may reduce power. The best performing approximate test, showing only mild deflation of P-values within the borderline genome-wide significance range (10^−5^ to 10^−6^), was an implementation of a Wald test, in which standardised log odds ratios under recessive and dominant models were tested against an *N*(0,1) distribution for significant departures from 0. In order to ensure null simulated P-values were close to their expected distribution at high levels of significance (about P<10^−4^), it was necessary to apply a standard correction to the estimated OR, adding 0.5 to each cell in the contingency table [37], and to account for this correction in the expected value of the test statistic when evaluating against the Null distribution. The best fitting inheritance model of the two (lowest P-value out of recessive and dominant) was taken as the P-value for the table in question, after multiplying by 2 to account for two tests being performed. Exclusion of variants with an observed minor allele frequency equal to or less than about 10% ensured accurate control of type-I error, and improved the appearance of volcano plots (effect sizes versus significance). (See SI and figure S8 for further details of these procedures.)

### Procedure for dealing with the relatedness between MZ and DZ twins

The MZ twins were dealt with by averaging over each member of the MZ pair and taking the average as a single individual, which reduces the influence of environmental variation in the associated facial phenotype. We dealt with the presence of DZ twins by performing 10,000 random selections of a list of unrelated DZ twins plus, for a random half of those individuals, their twin relative, as already described in the main text.

The most significant FDR-adjusted P-value [38] over the 10,000 selections was then taken as the overall observed P-value. Each random selection was again permuted to obtain 10,000 P-values distributed under the Null Hypothesis of no association. Related DZ twins had their genotypes drawn randomly according to Mendelian inheritance laws from 2 simulated parents (each with randomly drawn genotypes under Hardy-Weinberg Equilibrium using the observed allele frequency). Each related pair’s PC phenotypes were simulated as two N(0,1) distributed variables with 50% correlation. This is equivalent to the quite conservative assumption that the phenotype is 100% heritable. Then, individuals in the top theoretical decile of this distribution were designated as extremes before testing. Unrelated individuals were permuted as standard (random allocation of genotypes), and empirical P-values obtained by comparing the overall observed P-value with the permuted P-values, for each SNP separately (*P*=(*m* + 1)/(*n*+1), where *m* is the number of permuted P-values lower than the observed P-value and *n* the total number). These 2-tailed P-values were converted to 1-tailed values by dividing by 2 when the effect size was in the correct direction relative to the discovery analysis, and diving by 2 and subtracting from 1 when in the wrong direction.

### Frequency trend analysis

Combined TwinsUK and PoBI data were used to inspect for evidence for trends in genotype frequency across the quantitative PC phenotypes. This was done for each of the three replicating SNPs in the PC phenotype with which they were associated, in the PoBI data used for the discovery analysis. Individuals were sorted into 5% quantiles based on their ranked PC values. This was performed within the TwinsUK data using a randomisation scheme similar to that described above, in which a random set of unrelated DZ pair members was retained, and each time combined with the full set of phenotyped individuals from the PoBI discovery set, giving approximately 1467 individuals when analysing females only and 2118 individuals when analysing combined sexes. The mean frequency of genotypes in each bin for the combination of PoBI and TwinsUK data, the latter taken over the set of randomisations for DZ twins, was taken as the observed data for plotting.

### Function analysis of replicating SNPs

Using the TwinsUK sequence data (600 individuals), replicated variants were assessed for high *r*^*2*^ (as a measure of LD) with novel variants in the sequence data that were not present amongst the 524,576 SNPs used for the association analysis, and which were within 2Mb up- or downstream of the physical location of the replicated variant. Variants satisfying this requirement and with *r*^*2*^>0.2 were screened for functional information using SnpEff [39] and ensembl’s online tools (www.ensembl.org). Variants were determined to be putatively functional if they were a) exonic b) located within 5Kb up or downstream of a gene, c) within a UTR, d) within a splice site e) in a highly-conserved region, notably that could be coding for a functional RNA sequence or f) in a regulatory region, possibly a known enhancer. Inter-species conservation, another possible clue to function, was assessed using ensembl’s online tools by comparing the 10 base-pair flanking regions of variants among 8 primate species (Human, Chimpanzee, Gorilla, Orangutan, Green Monkey, Macaque, Olive Baboon and Marmoset).

Allele frequencies for replicating SNPs and LD-tagged putatively causal variants were assessed in several human populations from the 1000 genomes data [16].

## Acknowledgements

We thank the many researchers and volunteers who helped us at data collection events; in particular Ellen Royrvik, Helen Bodmer, Stephen Day and Ann Ganesan. We also thank the many volunteers who donated their 3D images and genetic data. This work was supported by Wellcome Trust grant 088262/Z/09/Z, EPSRC grant EP/N007743/1. The TwinsUK study was funded by the Wellcome Trust; European Community’s Seventh Framework Programme (FP7/2007-2013). The study also receives support from the National Institute for Health Research (NIHR)-funded BioResource, Clinical Research Facility and Biomedical Research Centre based at Guy’s and St Thomas’ NHS Foundation Trust in partnership with King’s College London. SNP Genotyping was performed by The Wellcome Trust Sanger Institute and National Eye Institute via NIH/CIDR

## Supporting Information

### Facial registration

The 3D face images generated by the 3dMD 3D camera system are provided in the form of a triangulated mesh (Fig. S1A). Each vertex has associated with it a 3D location and an RGB appearance value. However, the identity of each mesh vertex at this stage is unknown, and the number of vertices varies from image to image. Here we describe the process of fitting the mesh of a generic face model to this mesh; the result is a standardised triangulated mesh in which the identity of each node in the mesh is known (Fig. S1B). This process is known as mesh registration. A complete description of the process can be found in (Tena et al., 2006); more detail can be found in (Tena Rodriguez, 2007). We summarise the process here.

The generic face model consists of a triangulated mesh describing the surface of a human face. Each vertex of the model mesh has an initial 3D location and a predefined ID that identifies it as belonging to a specific part of the face. The model has an associated set of tools that can warp it to progressively align its surface with that of an input face mesh.

There are four main steps to the registration process:

1. Landmarking, in which 14 predefined salient landmarks are identified in the 3D input image. In order to match faces so that measurements at particular points correspond to one another in a meaningful way, each 3D photograph was manually ‘annotated’ at 14 landmarks. This involved placing a visual marker with a mouse cursor, for each photograph, on the nose tip, chin tip, labiomental crease, nasion, both corners of the mouth, the top and bottom of the lip, both sides of the nostrils, and each corner of each eye (Figure S1). This process was done manually by 3 different people for each photograph, and the mean position and corresponding model vertex ID recorded.
2. Global fitting, in which landmarks identified on the input mesh are used to warp the model so that corresponding landmarks in the model are brought into exact correspondence with them. The Thin Plate Spline algorithm (Bookstein, 1989) was used for the warping; this minimises the local deformation of the model as the landmarks are being brought into correspondence.
3. Local matching, in which, for each model vertex, the most similar vertex in the input mesh is identified. Similarity is defined here as the negative of the Euclidean distance between the vertices.
4. Energy minimisation, in which the model mesh surface is warped to align more closely with that of the input mesh, guided by the correspondences found in the previous stage. The energy E_tot_ that is minimised is the weighted sum of external and internal energy terms:

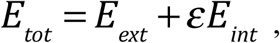

where ε is a weighting parameter, *E*_*tot*_ denotes the distance between the *n* input and model mesh vertices, {**x**_*i*_, *i* = 1…*n*} and 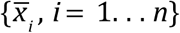respectively:

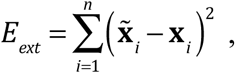

and *E*_*int*_ is a smoothness constraint that minimises the deformation of the model:

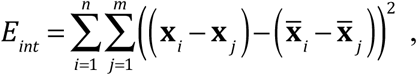

where ε is a weighting parameter 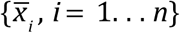 denotes the original positions of the model mesh vertices, and *j* = 1…*m* are the neighbours of vertex *i*. The weighting parameter *ε* was set to 0.25, and the conjugate gradient method was used for the energy minimisation.

Steps 3 and 4 of the above steps are combined in an iterative coarse-to-fine process. In the first two iterations, a reduced-resolution model (845 vertices) is used, while in the last two, the full-resolution model (3,300 vertices) is used. Hence the complete algorithm can be summarised as:

- Landmarking
- Global fitting
- Iterate 2 times with reduced resolution model:
  - Local matching
  - Energy minimization
- Iterate 2 times with full resolution model:
  - Local matching
  - Energy minimisation

### Heritability of registered data

Heritabilities of the registered face data were estimated using the 3dMD data collected from the TwinsUK sample described in the main text. For each facial variable in turn (total 3x29,658=88,974), the variables *V*_*MZ*_ and *V*_*DZ*_ were obtained by taking half the mean squared difference in phenotypic measurements between members of twin pairs, using the available 357 MZ and 394 DZ pairs. Heritabilities was then calculated as 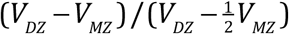 (Cavalli-Sforza and Bodmer, 1971). These were plotted as a heat map on the average face calculated from the PoBI data, assigning colours using the colorRampPalette function in R (Figure S2). The x variables are notable for having a strip of very low heritability down the center of the face, which is likely to be due to the situation of 6 landmarks in this region.

### Additive genetic value prediction (AGVP)

We present a method (denoted AGVP for additive genetic value prediction) for increasing the genetic signal present in a multidimensional phenotypic dataset where there are genetic correlations between measurements (i.e., where individual genetic variants, or perhaps multiple variants in linkage disequilibrium, affect a group of facial measurements). This is done without any reference to molecular genetic data. Rather, we interrogate the covariances between relatives’ (here, twins’) facial measurements. The resulting data, enriched for genetic signal, should be more amenable to subsequent genetic analysis.

The data consist of the raw facial surface measurements x_*ij*_, which are continuous random variables describing a position in one-dimensional space. *i* and *j* are indices for individuals (total *n*) and variables (total *m*) respectively. In the present application *m* ≈ 90,000, as there are approximately 30,000 registered ‘points’ each with positions in 3 dimensions).

To increase the heritability of facial measurements, we aim to estimate the unobserved values *E*_*e*_[x _*j*_ | *g*], where the expectation is taken over the stochastic environmental effects ( *E*_*e*_), for a given individual and for each measurement *j*, where x _*j*_ is the random variable of which x_*ij*_ is a realisation, and *g* represents a random vector of genotypes. Henceforth we denote *Y*_*j*_ = *E*_*e*_[x _*j*_ | *g*] as random variables with respect to genetic effects. In the quantitative genetics literature, the departure of *Y*_*j*_ from its population mean would be termed the *genetic value.* Under purely additive genetic effects (no dominance), the assumption we rely on below, it is known as the *breeding value* or *additive genetic value*.

The objective is to predict *y*_*ij*_, the realised additive genetic value for individual *i*, which cannot be directly observed, using the set of measurements taken on the same individual: {x_*ik*_ :*k* ∈1,2,3…*m*}. This is performed for each *j* in turn. Each x_*k*_ is modelled as

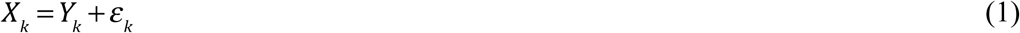

for all *k* in 1,2,3…*m*, where *ε*_*k*_represents the effects of the environment. We assume that *E*_*e*_[*ε*_*k*_| *g*]= 0, i.e. that there are no gene-environment interactions.

We minimise the expected least squares error

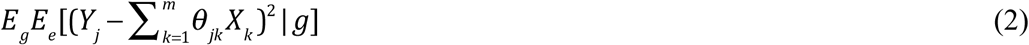

with respect to each coefficient θ _*jk*_, which represents the predictive influence of variable x_*k*_ on variable x _*j*_ This double expectation is taken with respect to the stochastic environmental and genetic effects over individuals ( *E*_*e*_ and *E*_*g*_ respectively).

#### Unbiasedness Constraint

We apply the constraint

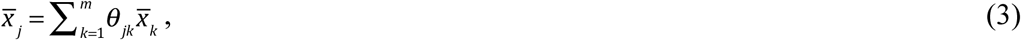

where *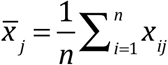*. Put another way, the predictor, when run on the ‘average *n*individual’, must return the values for the ‘average individual’. Under the proposed model (Expression 1) *E*_*g*_*E*_*e*_[x_*k*_ | *g*]= *E*_*g*_*E*_*e*_[*Y*_*k*_ | *g*], so, asymptotically, the constraint implies that

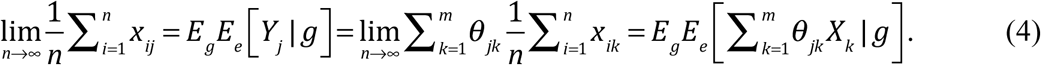

However, without applying this constraint, *E*_*g*_*E*_*e*_[*Y*_*j*_ | *g*] does not necessarily equal 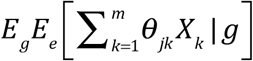 and so the estimator in Expression 2 is not necessarily unbiased.

With the constraint applied, the quantity to be minimised becomes

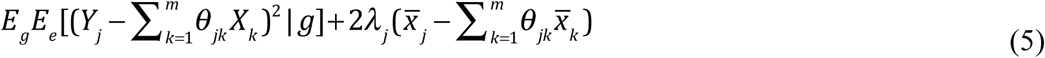

where λ_*j*_ is a Lagrangian parameter.

#### Solution using additive genetic covariance

As it is assumed that the expectation of the environmental effects, conditioned on genotype, is zero (i.e. that there are no gene-environment interactions) then, using Expression 1,

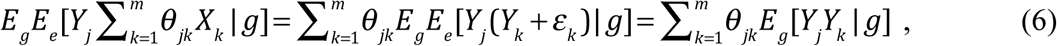

so, Expression 5 expands to

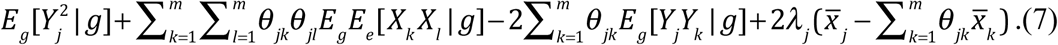

After cancellation of terms that are asymptotically equal due to the Lagrangian constraint, Expression 7 becomes

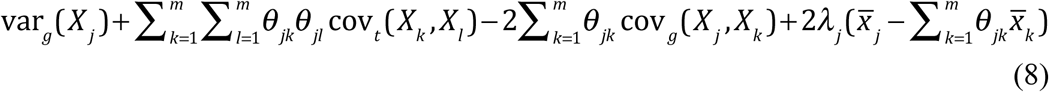

where cov_*t*_ (x_*k*_, x_*l*_) represents the total covariance between variables *k* and *l,* var_*g*_(x _*j*_) represents the genetic variance component for variable *j* and cov _*g*_(x _*j*_, x_*k*_) represents the genetic covariance component between variables *j* and *k*. To proceed, we assume that the full effects of alleles can be well-approximated by their additive effects, and so substitute cov _*g*_(x _*j*_, x_*k*_) with cov_*a*_(x _*j*_, x_*k*_), the additive genetic covariance, which can be estimated by taking the covariance between twins with respect to the population mean, devalued by inverse relatedness:

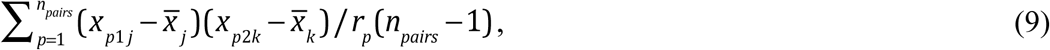

where *p* is a twin pair index, *n*_*pairs*_ is the number of pairs, x_*p*1 *j*_ is the measurement on the first of the pair for variable *j*, and x_*p2k*_ the measurement on the second of the pair for variable *k. r*_*p*_ is the coefficient of relationship between the twins in the pair: 1 for monozygotic (identical) twins and 0.5 for dizygous (non-identical) twins. Expression 9 assumes that effects attributable to pairs’ shared family environments are negligible, though it would be possible to accommodate for this using standard techniques (Falconer and Mackay, 1996). Shared family environment components are typically found to be very small (Plomin, 2011), so we take this assumption to be reasonable. It is possible, using Expression 9, to estimate the additive genetic covariance using pairs of more distantly related individuals, e.g. those from a population sample, so long as sufficient molecular genetic data are available for calculating their coefficients of relationship.

To minimise Expression 8 we differentiate with respect to each θ _*jk*_. Setting the resulting partial derivatives zero, we obtain the solution

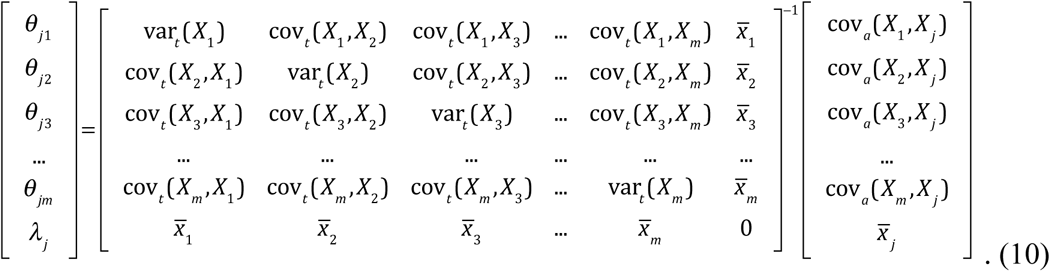

The predicted additive genetic values, for each *i* and *j*, are then 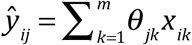. The estimator in Equation 10 is analogous to the *Universal Kriging* estimator used to predict quantities for variables of interest, e.g. mineral levels, at particular geographic locations, based on measurements of the same variable taken at other locations nearby (Cressie, 1991). The variance of *ý* _*ij*_ is 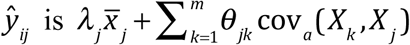, though this does not account for sampling error affecting the estimates of variances, covariances and sample means, (i.e. it assumes that those estimates equal their true values).

As each variable *j* is treated independently, all of their coefficients can be represented in the single equation **P** = **T**^−1^**A**, where

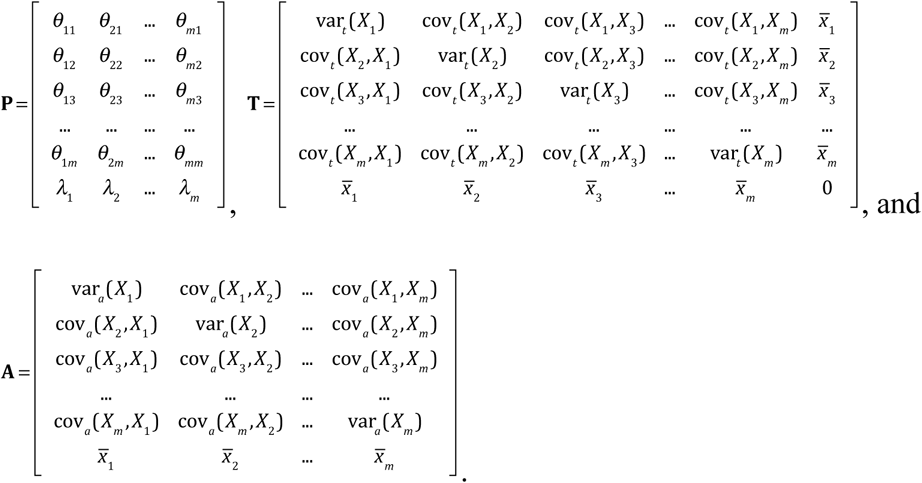

#### Fitting process using facial subsets

Ideally the linear predictor uses all facial variables {x_*ik*_ :*k* ∈1,2,3…*m*} as described above, where *m* is the total number processed and registered from the camera system (approximately 3x30,000=90,000). However, computational limitations prevent this, so a workaround is proposed whereby analysis is restricted to rectangular subregions of the face. These are selected based on visual inspection of an image of the average face (Figure 1). Vertices that fall within the subregion on the average face are taken forward for AGVP analysis in all individuals, so that the number of variables remains constant.

Two subregions, henceforth referred to as ‘profile’ and ‘eyes’, were defined for AGVP analysis and subsequent genetic association mapping. The constituent vertices of these sub-regions are highlighted according to their positions on the average face in Figure 1. These two particular regions were chosen, fairly informally, on the basis that they among the most strongly identifying aspects of the face. The eyes subregion contains 2763 vertices, each in 3 dimensions, giving 3 x 2763 = 8289 variables for analysis, whereas the profile subregion contains 1646 vertices. Only the Y (height) and Z (depth) dimensions were used, giving 2 x 1646 = 3292 variables for analysis, as the x (width) dimension of the profile is both less characteristic of appearance and less heritable (Figure S2A).

#### 5-fold cross validation for protection against overfitting

The method above is susceptible to overfitting due to the involvement of a large number of variables. To guard against this, a ridge penalty is applied to the parameters {θ :*k* ∈1,2,3…*m*} so that **T^−1^**is now

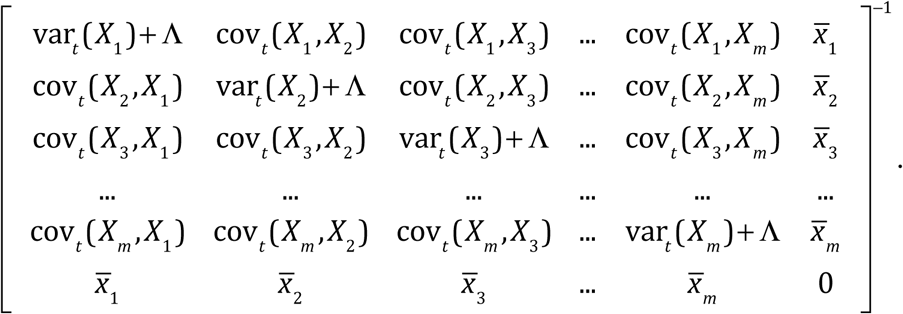

Predicted additive genetic values cannot be compared with any measured values in order to calculate an error rate for the fitted model, as it is not possible to observe a additive genetic value directly. However, additive genetic values, representing purer genetic effects rather than raw phenotypes, should have higher heritabilities than their associated raw measurements. Therefore, 5-fold cross-validation was used to find a value of Λ that gave high-quality additive genetic value predictions as measured by the mean heritability taken across variables. 1567 TwinsUK individuals were split into 5 validation sets, each consisting of 50 MZ and 50 DZ pairs (200 individuals), and a test set consisting of 107 MZ and 144 DZ pairs (502 individuals). Leaving out a single validation set, and for a fixed value of Λ, AGVP analysis was performed using the remaining 865 individuals (200 MZ and 200 DZ pairs, plus 65 unrelated individuals), and the fitted model applied to each validation set in turn. Using the excluded validation set alone, heritability was calculated for the predicted additive genetic values, and the mean heritability taken across all facial variables under analysis. This was repeated in turn for each validation set, and the mean and standard error of the heritability score taken over the 5 sets, giving an overall score with associated standard error for the starting value of Λ. This process was repeated for different values of Λ, using identical validation and test sets, thus obtaining a mean and standard error for the heritability score for each Λ. The range values over which to increment Λ was set to 1,2,3…10, for both subregions, based on experimentation with heritability results from AGVP analysis. Using the ‘one standard error’ rule, the optimal value of Λ was chosen by subtracting one standard error from each mean heritability score, and picking the Λ giving the highest value. For both the profile and eyes subregions, the optimal value was Λ = 10. Finally, this optimal value was used to fit a AGVP model using all the data apart from the test set, which was used to assess the heritabilities in the same way, taking heritabilities of predicted additive genetic values for each phenotypic variable, then taking the mean across variables. The mean heritability was 76.1% for the eyes sub-region and 81.5% for the profile sub-region, compared to 69.8% and 76.6% for the original variables respectively. Perhaps more importantly, the variance in heritabilities was also reduced, as variables with very low raw heritabilities had much improved heritabilities in the predicted additive genetic values (Figures 2 and S3).

**Fig. 2.**
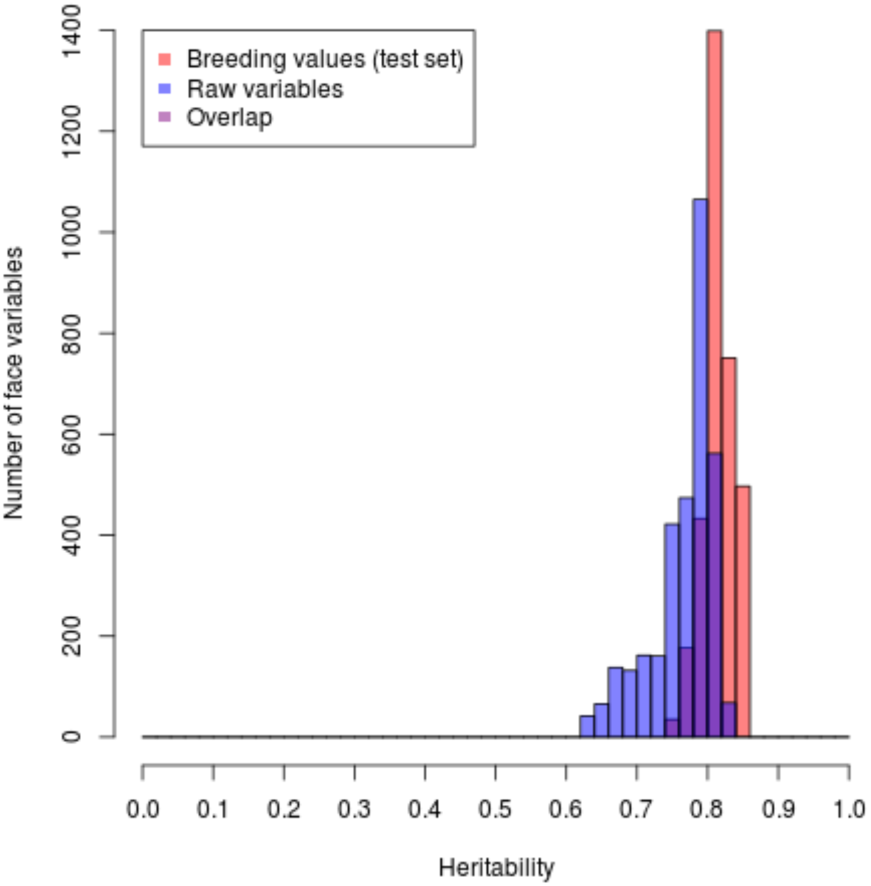
Comparison of profile heritabilities for raw variables versus additive genetic values.

### Selecting a subset of facial Principle Components

Shape translation and PCA were performed as described in the main text. The largest 50 PC axes then were inspected for promising phenotypes using two criteria:

a) Heritability of each PC axis as determined using the PC scores of the 1567 TwinsUK individuals and b) The squared difference between the mean PoBI (European) PC score and the mean East Asian score, taken as a ratio against the within-population variance. Plots of these statistics are shown in Figure S4. PCs with heritability>0.75 or heritability>0.65 and a population variance ratio of >1 were taken forward for genetic association analysis.

### Genotype quality control

#### Discovery data

The intersection between the two platforms (Illumina Human 1.2M-Duo and Illumina Infinium OmniExpress-24 BeadChip 750K) was 547,863 SNPs. Prior to QC there were 3,735 genotyped DNA samples, constituting 3,616 unique individuals.

151 individuals were removed due to unusual genotyping on the sex chromosomes that did not correspond with their reported sex. Self-reported females were removed if Y chromosome missingness was less than 50% and excess homozygosity on the x chromosome, as determined using plink’s F-statistic, was greater than 0.3. Similarly, self-reported males were removed if Y chromosome missingness was greater than 50% and excess homozygosity on the x chromosome was less than 0.8. 14,987 SNPs and 63 individuals with genotyping rates less than 1% were discarded. 52 individuals were removed for having a genome-wide F statistic greater than 3 standard deviations from the population mean; all 52 showing an excess rather than deficit in homozygosity, probably due to parental relatedness. One individual with an extreme F statistic of 0.13 was confirmed from our paperwork as having parents who were probably first cousins. Markers were removed due to showing evidence for a departure from Hardy-Weinberg Equilibrium (3276 SNPs), using a cutoff of P less than or equal to 10^−4^. 24 individuals showed some remaining evidence of a mismatch between self-reported and SNP-determined sex (based on plink’s x-chromosome homozygosity scores with F<0.2 suggesting female and F>0.8 male) and were removed due to probable mislabelling or contamination. Of these, 21 were in the WTCCC2 exclusion list (Genetic Analysis of Psoriasis et al., 2010) and 3 were genotyped after the WTCCC2 study. 5 samples’ sex discrepancies were best explained by ID mismatches, as they showed full relatedness with other individuals. Relatedness was assessed using a subset of 74,615 SNPs with pairwise r^2^<0.1. A number of samples otherwise passing QC were removed due to close relatedness or due to being duplicates of the same individual. 64 known duplicated samples and 187 individuals with an identity-by-descent statistic greater than 0.125 (equivalent to first cousins) with at least one other individual were removed. 30 of these samples were removed due to full relatedness with a different individual’s sample. All apart from 2 could be attributed to either a) genotyping, contamination or misidentification issues meaning that they were present in the WTCCC2 exclusion list, b) multiple sample collection events visited by the same person, identified by examining paperwork, or c)in one instance, identical twins.

Principal Components Analysis was performed using the 74,615 remaining SNPs with r^2^<0.1 and the largest 5 axes inspected visually for outlying samples. The largest Principal Component contained a cluster of 33 outlying individuals (> 3 standard deviations from the mean) that could not be attributed to population structure or batch effects. A large proportion of these individuals were found to be nearby to one another on the same genotyping plate, suggesting contamination of DNA. All 33 samples were removed.

As genotyping was performed in 7 different batches, 2x3 chi-square tests were performed in turn between each batch and all other combined batches, for each SNP, to detect possible batch effects. Using a P-value threshold of 10^−6^, 5024 SNPs were found significantly associated with genotyping batch, and were consequently removed. After QC, genotypes for 3161 individuals (1532 male, 1629 female) and 524,576 SNPs were retained for association analysis.

#### Replication data

TwinsUK genotype data was available on two platforms; 1278 samples represented by 2,287,998 array-typed SNPs and 612 whole-genome sequenced samples (19,725,734 autosomal variants). Separate QC procedures were performed for the two platforms before merging. For the array data, 81 samples with a genotype call rate less than 5% were removed. Examining the distribution of F-statistics for individuals with extreme levels of homozygosity, 3 individuals stood out as clear outliers with F statistics 4 standard deviations above the mean, and were removed. For sequenced samples, 326,065 variants with genotyping call rates of less than 1% were removed. Distributions of heterozygosity F-statistics were examined and 12 outlying individuals greater than 3.25 standard deviations from the mean removed (7 with excess and 5 with deficient heterozygosity). After merging the array and sequencing data and retaining variants common to both sets, 1,887,250 variants and 1794 samples, 1275 of whom were unique individuals. Duplicate samples due to individuals being both array-typed and sequenced were eliminated by discarding the array sample. The largest PC axes derived from the genotype data were inspected for any evidence of batch effects between array-typed and sequence data, with none found.

### Candidate SNP list

Candidate genes were chosen by reviewing the literature on human facial dysmorphias and facial morphologies in other species, in order to produce a list of regions with prior evidence for involvement in facial features, that can be therefore be subjected to less-stringent multiple comparisons correction. Based on the literature findings, additional searches were performed in the Online Mendelian Inheritance in Man (OMIM) and Ensembl databases. There are 381 genes/regions in the candidate database, representing 168 different dysmorphic facial features. Large candidate regions (approximately 1Mb or larger) were broken down so that only the genic subregions were retained. SNPs were designated candidates if they were located within 75Kb of a candidate region, allowing for LD tagging of functional variants by non-functional SNPs physically close to the relevant gene.

### Genetic association analysis (discovery)

In order to investigate the hypothesis that there are genetic variants conferring large effects on facial morphology, we focus on individuals with facial features that can, in some sense, be considered ‘extreme’ relative to the general population. We therefore dichotomised our PCA based facial phenotypes into subsets of upper and lower extremes, which constitute the top and bottom 10% of individuals when ranked according to their scores for each PC. Ranking was performed within 1423 individuals (652 males and 771 females) for whom both genotype and phenotypic data were available. Ranking was performed separately within each sex. For visualisation purposes, average faces within each extreme were produced using a Matlab script, by plotting the arithmetic mean for each coordinate measurement in each vertex, among all individuals falling into the designated extreme, and overlaid with a surface texture. Only females were used to produce average faces, so as to facilitate comparison with the TwinsUK data, which are almost entirely female. Average faces were also produced using all Chinese females and all British (PoBI) females separately.

For each PC in turn (5 in each subregion), upper extremes were tested against all remaining individuals (including both the lower extremes plus the 1738 not phenotyped PoBI individuals remaining after genotype QC) by analysing the 3x2 tables of genotypes (*aa/Aa/AA* where *a* represents the minor allele) versus extreme/control status, for all 512,181 autosomal SNPs. This procedure was repeated for the lower extremes, combining not phenotyped PoBI individuals with upper extremes as the control sample. All association analyses were performed 3 times; for female extremes (excluding male phenotyped individuals - approx. 77 extremes, 2478 controls), male extremes (excluding female phenotyped individuals - approx. 65 extremes, 2377 controls) and combined male and female extremes (approx. 142 extremes, 3019 controls). In total, there were 6 association analyses performed for each PC under consideration, due to separate testing of both upper and lower extremes.

The relatively small number of extreme individuals motivated a careful selection of the appropriate statistical test for contingency table significance. This is especially pertinent when examining models of inheritance involving the effects of minor allele homozygotes, which can be quite rare even when an allele is common. In order to establish the most appropriate test, null hypothesis simulations of 3x2 tables were performed, and the type-I error rates quantified. Previous work (Bigdeli et al., 2014) has demonstrated that, when the numbers of observations are small and very low P-values are required, e.g. less than 5x10^−8^ for genomewide significance, a variety of traditional frequentist tests are conservative, either due to asymptotic assumptions breaking down, or due to inherent conservativeness in the case of Fisher’s Exact test. Simulations performed under our study design, which in contrast with Bigedli et al., had unequal proportions of extremes versus controls, found that most of these tests are anti-conservative. Fisher’s exact test controlled the type-I error well in simulations, but was not chosen as it is often conservative and may reduce power. The best performing approximate test, showing only mild deflation of P-values within the borderline genome-wide significance range (10^−5^ to 10^−6^) (Figure S5), was an implementation of a Wald test, in which standardised log odds ratios under recessive and dominant models were tested against an *N*(0,1) distribution for significant departures from 0. In order to ensure null simulated P-values were close to their expected distribution at high levels of significance (approx. P<10^−4^), it was necessary to apply a standard correction to the estimated OR, adding 0.5 to each cell in the contingency table (Haldane, 1956), and to account for this transformation in the expected value of the test statistic when evaluating against the Null distribution (Details in caption of Figure S5). The best fitting inheritance model of the two (lowest P-value out of recessive and dominant) was taken as the P-value for the table in question, after multiplying by 2 (up to a maximum value of 1) to account for two tests being performed. Variants with an observed minor allele frequency of less than 10% were excluded to further ensure accurate control of type-I error, and to improve the appearance of volcano plots (effect sizes versus significance).

Distribution of 10^8^ P-values from Pearson’s chi-square test, under the same Null hypothesis simulations, showing serious deflation. Similar results are obtained when using Yates’ correction.

The Wald test was applied to the PoBI extreme/control PC-based phenotypes for all SNPs passing QC. Figures 3, S6 and S7 display results from the discovery analysis relating to SNPs that subsequently replicated in our follow up analysis. Figure S5B shows the serious departure from the expected linear relationship between the observed and expected P-value quantiles that is obtained for the Pearson’s Chi square test as compared to the Wald test.

**Fig. 3.**
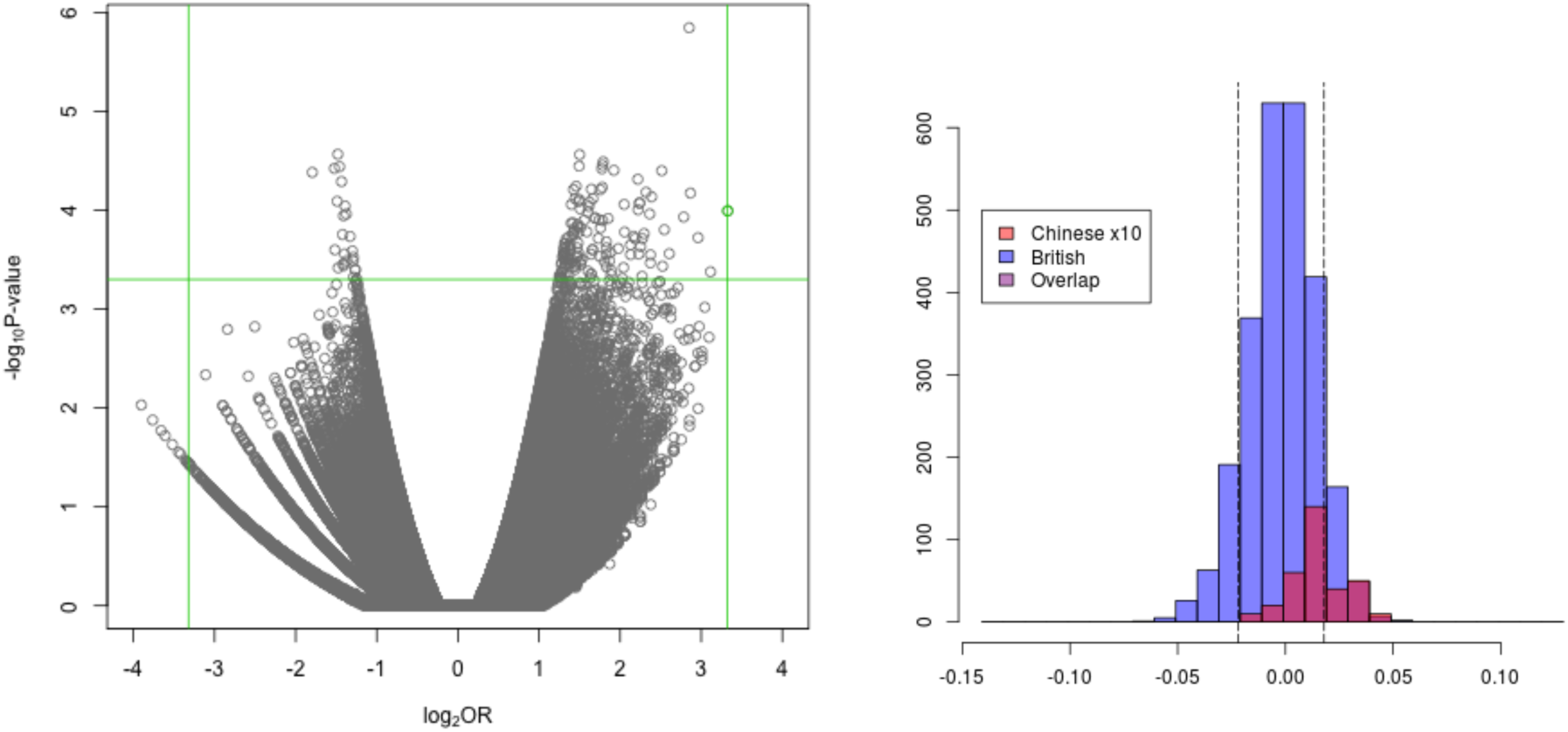
(A) Volcano plot for females lower (more European) extreme PC2 profile.

### Replication analysis

Variants passing any of the three criteria for follow-up were analysed in the TwinsUK cohort (see the ‘genotype quality control’ section for QC procedures). Although this dataset was used previously to produce estimates of the additive genetic values, which were the foundation of the discovery analysis above, it nevertheless constitutes a valid and unbiased replication set, as the AGVP method makes no reference to DNA data.

1271 of the 1275 TwinsUK individuals with genotype data after QC had available facial phenotypic data, 599 of which were sequenced and 1190 array-typed (with 518 being both sequenced and array-typed). One complication of replication analysis was the presence of many related individuals in the TwinUKs data. Among the 1271 with both genetic and phenotypic data, 536 were MZ twins and 734 were DZ twins, with one individual of unknown twin status and no relations in the database. The median age was 61 (mean=59.50, sd=9.70) upon the date of photographic phenotyping (between 2009-2012 with most photographed in 2010/2011). Just 4 of these individuals were male.

As in the discovery analysis, individuals were ranked according to their PC scores and categorised into upper and lower extremes (top and bottom 10%). Average faces were calculated, within each extreme, as in in the discovery analysis. All profile- and eyes- associated discovery SNPs were tested for association in the replication dataset using Wald tests as in the discovery analysis. A notable difference between the two analyses is that almost all of the TwinsUK samples have phenotypic data available, whereas 1738 of 3161 PoBI individuals were not phenotyped and treated as ‘unscreened’ control samples. The correction for discrete distributions (adding 0.5 to each contingency table cell) was not applied to the replication Wald tests, as permutation was used to ensure accurate control of Type-I error, and was computationally feasible due to the relatively small number of required tests.

In total, 13 profile-associated and 6 eyes-associated SNPs were taken forward for replication analysis. Of the 7 eye-associated discovery SNPs, 1 (rs2039473, eyes PC3 associated in females) was not present on the TwinsUK genotyping platform and not pursued further. After removing a single SNP having r2 > 0.1 with another discovery hit (retaining the SNP with the highest OR in the discovery analysis), 12 remained associated with profile phenotypes, with all 6 remaining eyes-associated SNPs retained as independent associations. For each of the 18 SNPs, association was only tested for the extreme, either upper or lower, and PC with which it was associated in the discovery analysis. To increase power, control samples from the appropriate discovery analyses were incorporated into the replication contingency tables before testing. For each SNP, a Wald test was performed under whichever inheritance model was found most significant in the discovery analysis (recessive in all cases), and the total number of tests performed was 18, equal to the number of SNPs.

A complication in using the TwinsUK cohort as a replication dataset is the high degree of relatedness between MZ and DZ twins. Removing all related individuals satisfies the requirements for independent samples required by standard statistical analyses, but also loses information and results in an arbitrarily chosen set of unrelated samples. To address these issues, we first treated MZ twins as a single composite individual by averaging over their phenotypic measurements; for each PC phenotype (before assignment of Extreme/Control status) we took the mean of each MZ pair and assigned this phenotypic value to their shared genotype (choosing a sequenced member of the pair over an array-typed member wherever possible). Averaging over each member of the MZ pair reduces the influence of environmental variation in the facial phenotype associated. After removing one member of each MZ pair, leaving a sequenced over an array-typed member where possible, 1029 individuals remained (1027 female and 2 male). 737 were DZ individuals (327 pairs plus 83 with no remaining related twin) and 291 MZ twins remained without any of their related pair members. One individual had unknown twinning status, with no related individuals present in the database.

We dealt with the presence of DZ twins by performing 10,000 random selections of a list of unrelated DZ twins plus, for a random half of those individuals, their twin relative. The most significant FDR-adjusted P-value over the 10,000 selections was taken as the overall observed P-value. Each random selection was again permuted to obtain 10,000 P-values distributed under the Null Hypothesis of no association. Related DZ twins had their genotypes drawn randomly according to Mendelian inheritance laws from 2 simulated parents (each with randomly drawn genotypes under Hardy-Weinberg Equilibrium using the observed allele frequency). Each related pair’s PC phenotypes were simulated as two N(0,1) distributed variables with 50% correlation. This is equivalent to the quite conservative assumption that the phenotype is 100% heritable. Then, individuals in the top theoretical decile of this distribution were designated as extremes before testing. Unrelated individuals were permuted by randomly shuffling their case/control statuses. Empirical P-values were once more obtained by comparing the overall observed P-value with the permuted P-values. For each SNP separately, the observed P-values were adjusted for 10,000 tests being performed using the Benjamini-Hochberg method (Benjamini and Hochberg, 1995), and the most significant P-value after adjustment taken as the overall single P-value for that SNP. This was then compared with the 10,000 permutated P-values to obtain an empirical P-value (*P* = (*m* + 1)/(*n* + 1) where *m* is the number of permuted

P-values lower than the observed P-value and *n* the total number). These were converted to 1-tail tests by dividing by 2 if the effect size was in the correct direction (based on the expectation from the discovery analysis) or dividing by 2 and subtracting from 1 otherwise. The distribution of these empirical, 1-tailed P-values, as compared to the expected values based on their ranks, is shown in Figure S8, with those passing an FDR of 5% highlighted, and the deviation from the expected line under no associations is clear.

### Analysis of combined discovery and replication data

Contingency tables for replication analyses were produced using the rounded mean cell counts over random selections, and the median OR over random selections was taken as an estimate of the OR. The combination of these tables together with the PoBI discovery data was used to perform a Wald test (applying the 0.5 correction factor) for significance and estimate the overall OR.

### Assessment of evolutionary conservation

Species comparisons were made between humans and primates using ensembl’s online tools (www.ensembl.org), shown in Figure S9.

#### Genetic analysis of extremes in TwinsUK sequence data

Variants that were identified as putatively causal were further analysed in sequenced members of the TwinsUK cohort (n=600). Dominant or recessive ORs for putatively functional SNPs that had similarly high OR as the discovery SNP were taken to be further evidence for the causal involvement of the putatively functional variant in the phenotype. Dominant or recessive ORs were calculated using extreme and control statuses assigned in the same way as before, using the reduced set of 600 individuals.

---

**Fig. 6.**
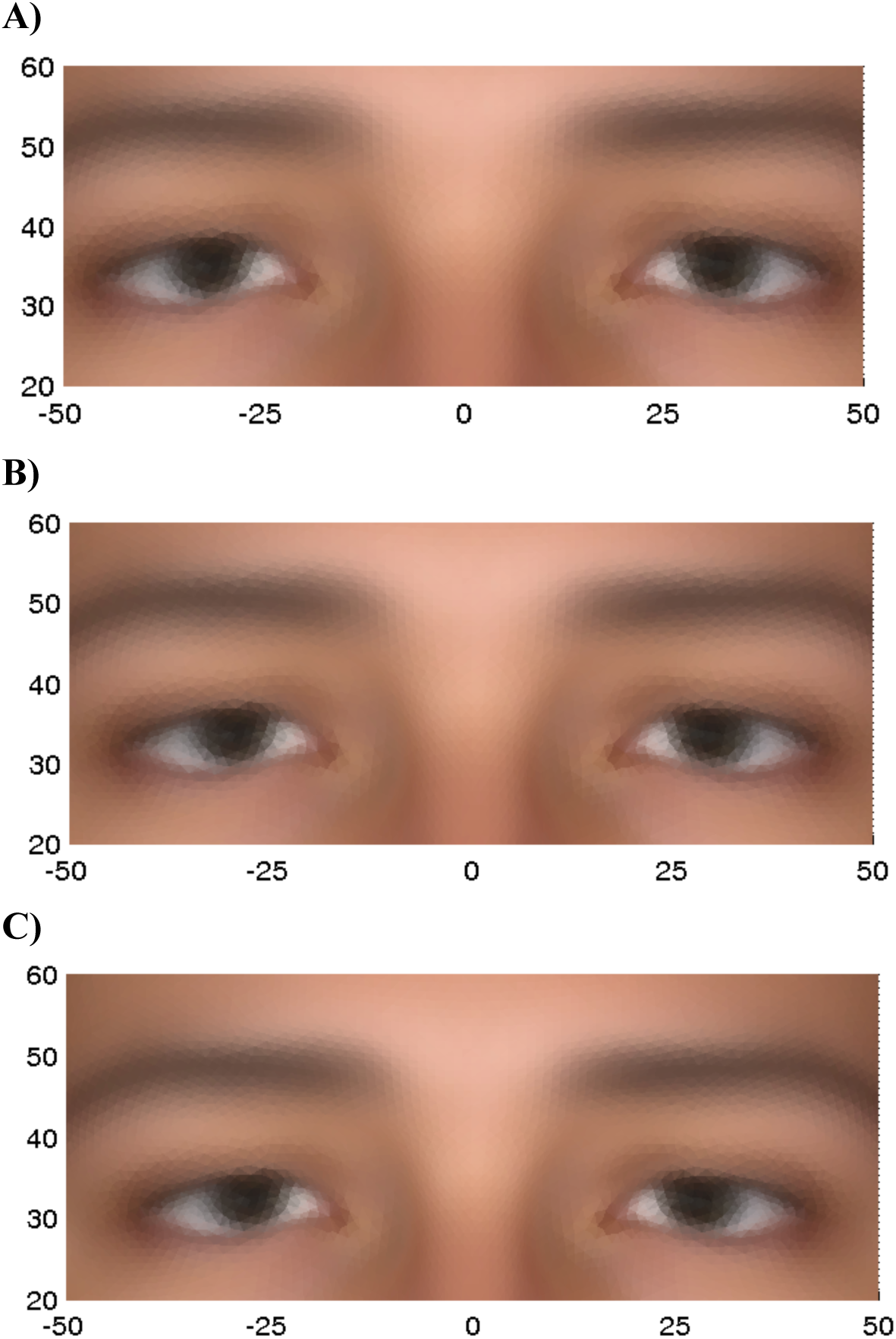
Average eyes phenotype for PC1 (females only) for the upper(A) and lower (C) extremes and the overall average(B)

**Fig. S1.**
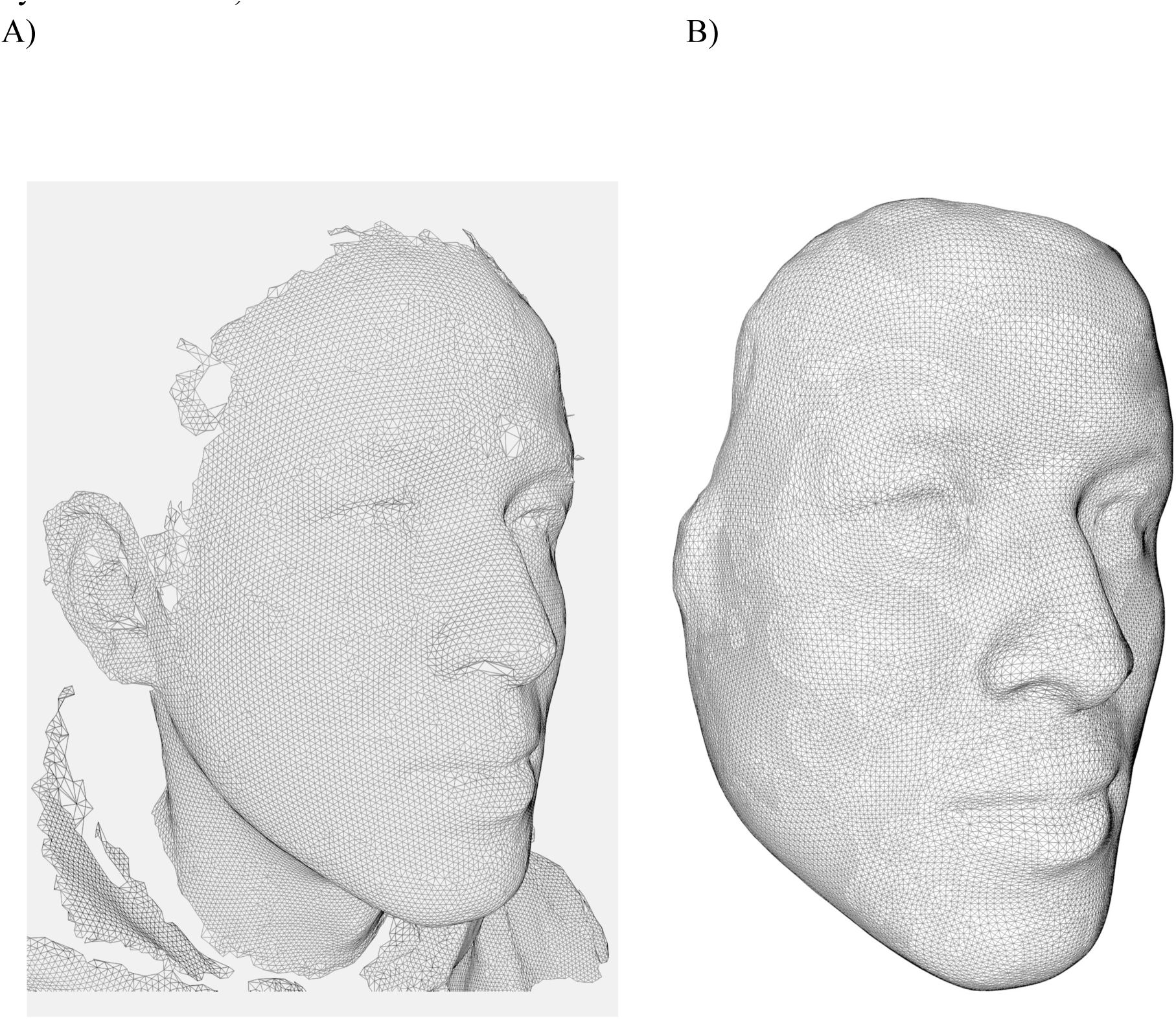
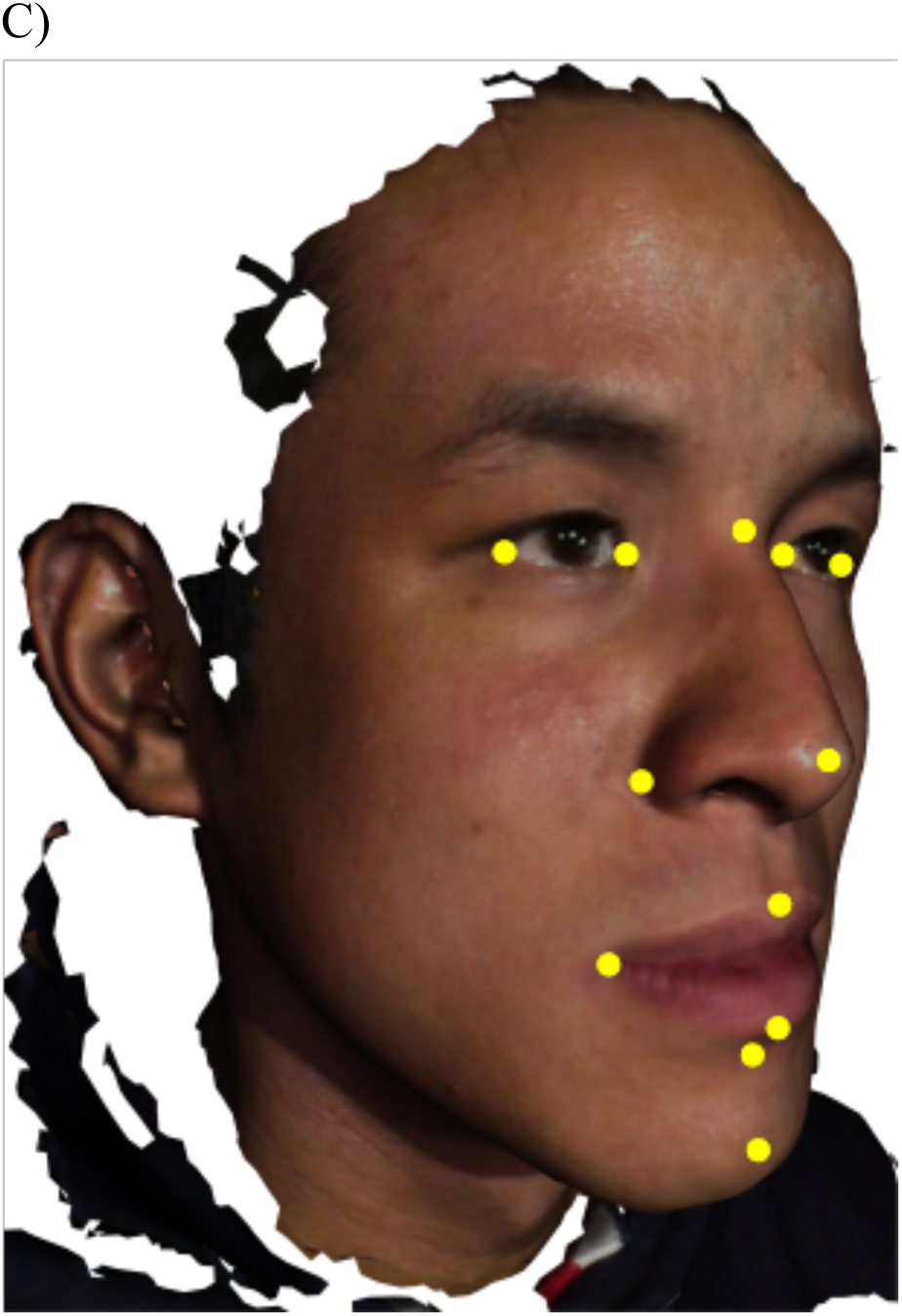
Example of triangulated mesh prior to (A) and after (B) registration, and locations of 14 manual landmarks (C) (with left corner of mouth and left corner of nose obscured by the orientation).

**Fig. S2.**
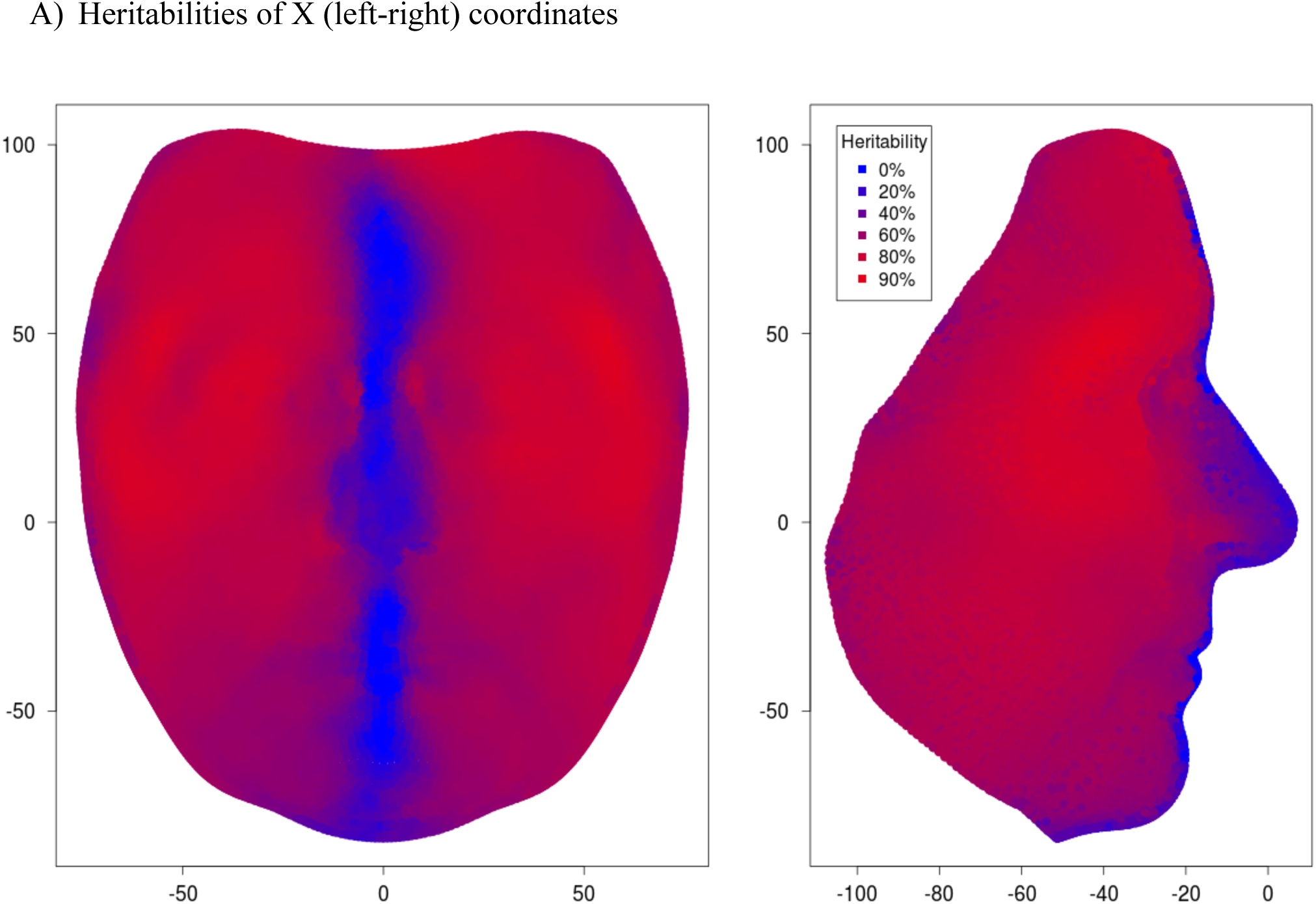
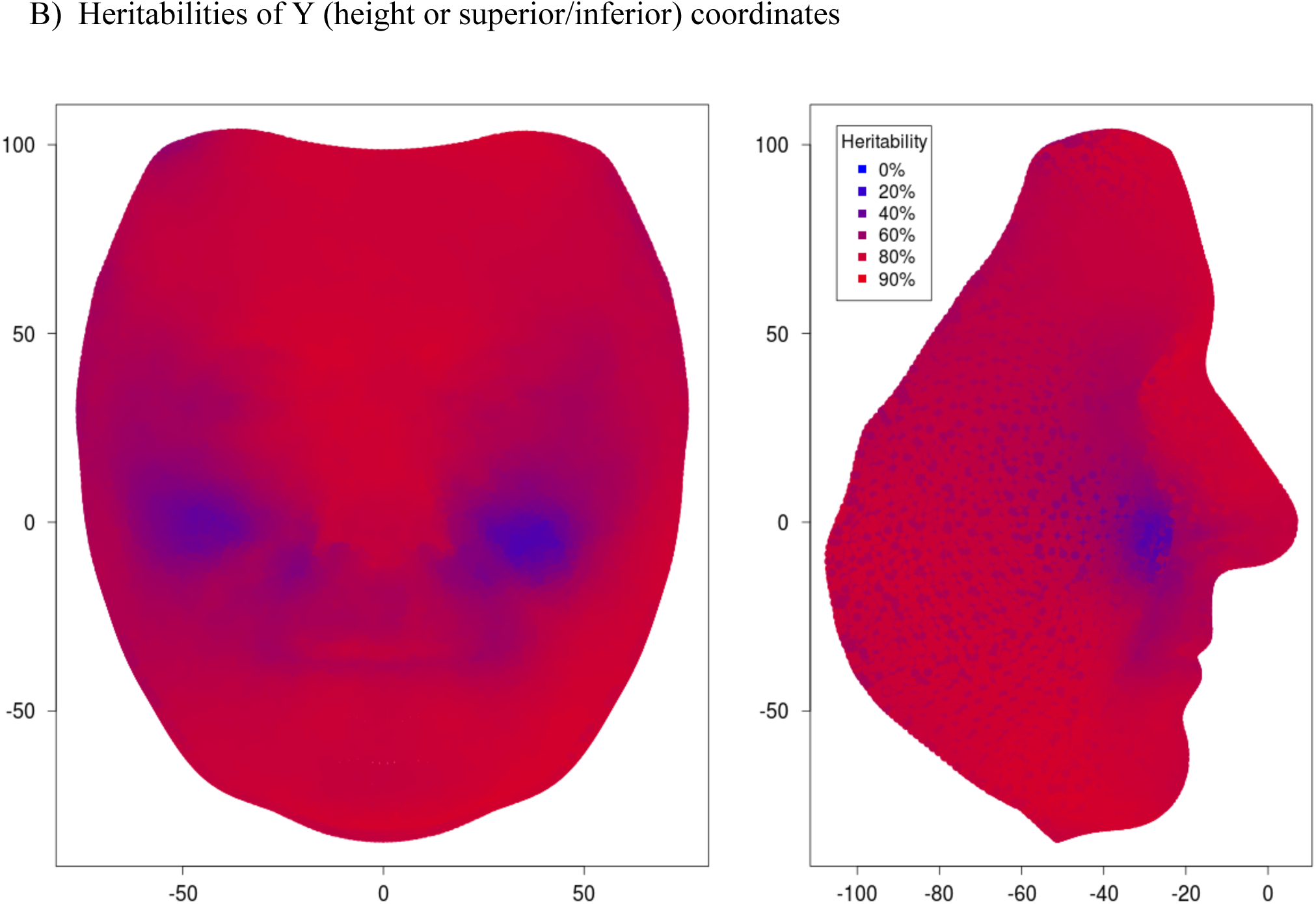
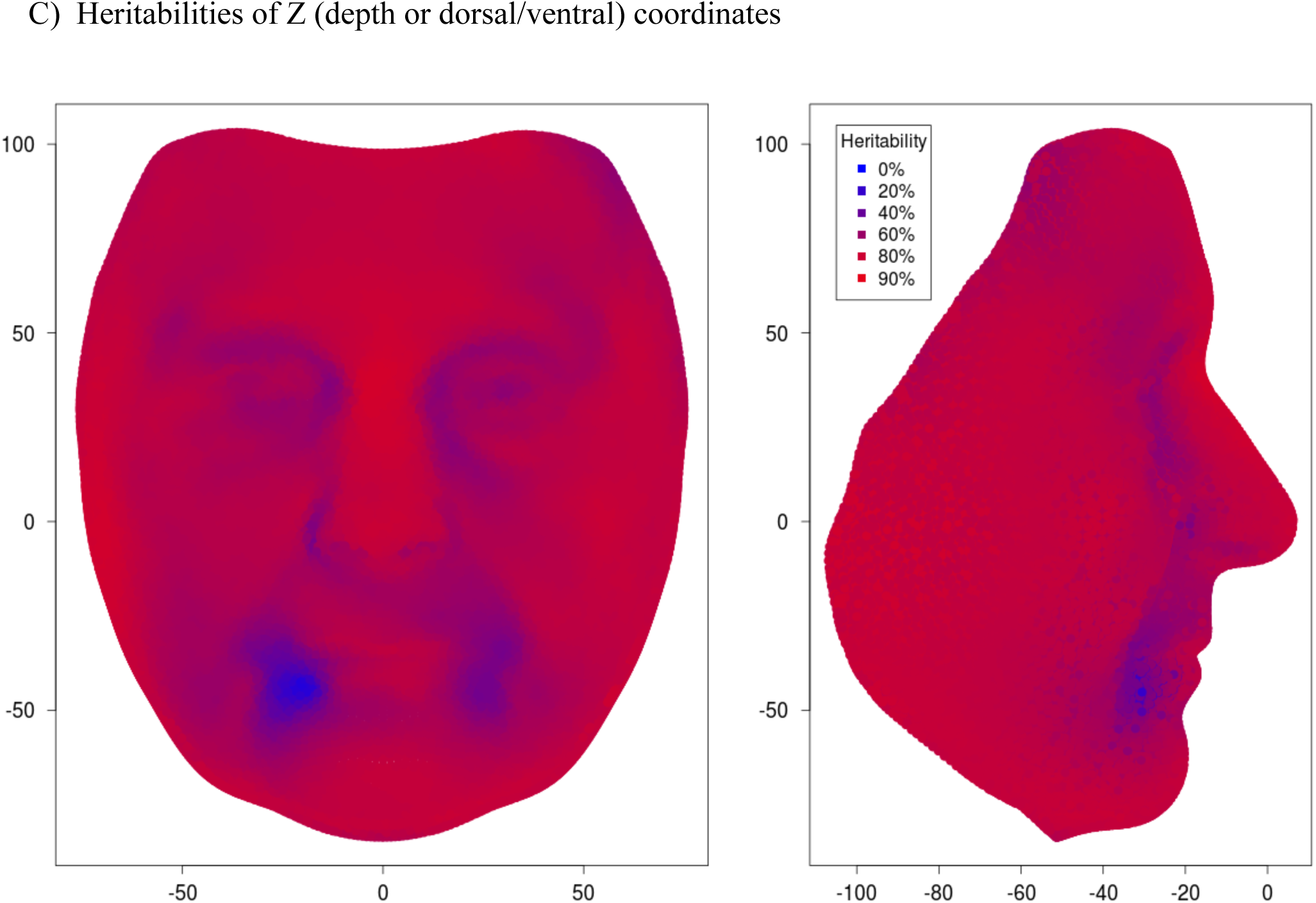
Heritabilities of ‘raw’ vertex data, after registration but before transformation into additive genetic values, plotted on the average face.

**Fig. S3.**
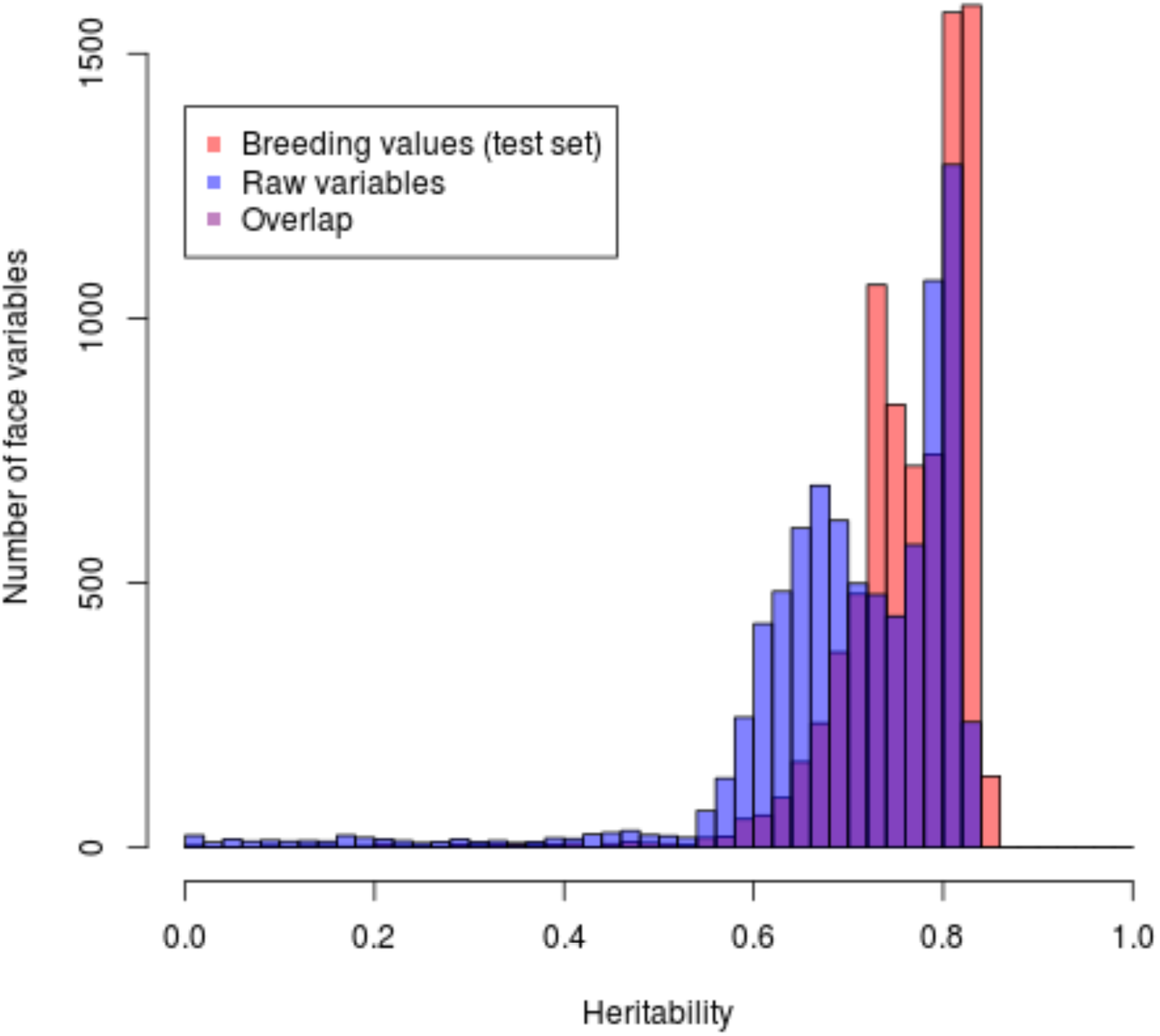
Comparison of heritabilities in raw variables versus additive genetic values (calculated using an independent test set) for the eyes subregion. Figure 2 in the main text shows the equivalent data for the profile subregion.

**Fig. S4.**
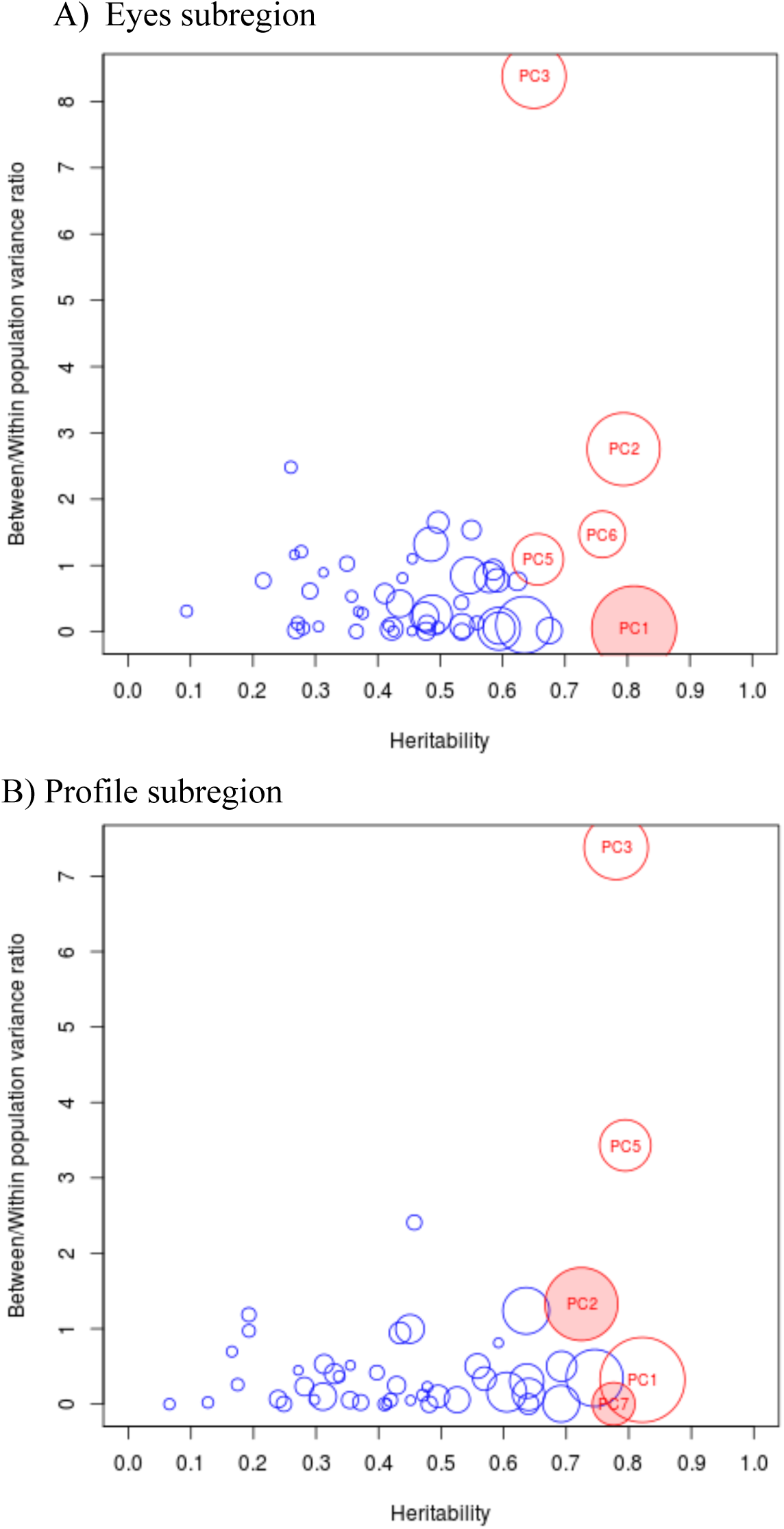
European/East-Asian differences (between/within population variance ratios) and heritabilities for largest 50 PCs performed on facial additive genetic values. Size of circles indicate the rank of the corresponding PC, and red circles denote PCs taken forward for genetic association analysis. Filled in circles denote phenotypes for which SNP associations were replicated. The x-axis has been limited to the interval [0,1]. Some small PCs had heritabilities below zero.

**Fig. S5.**
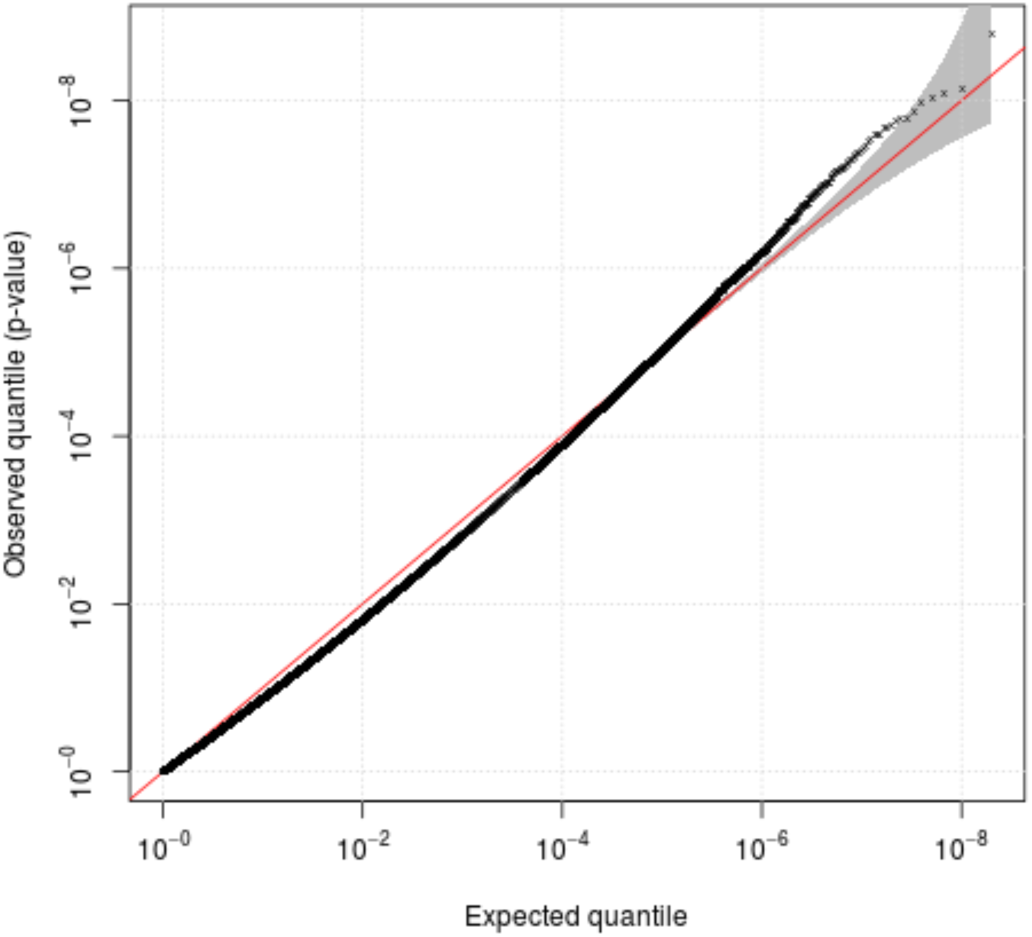
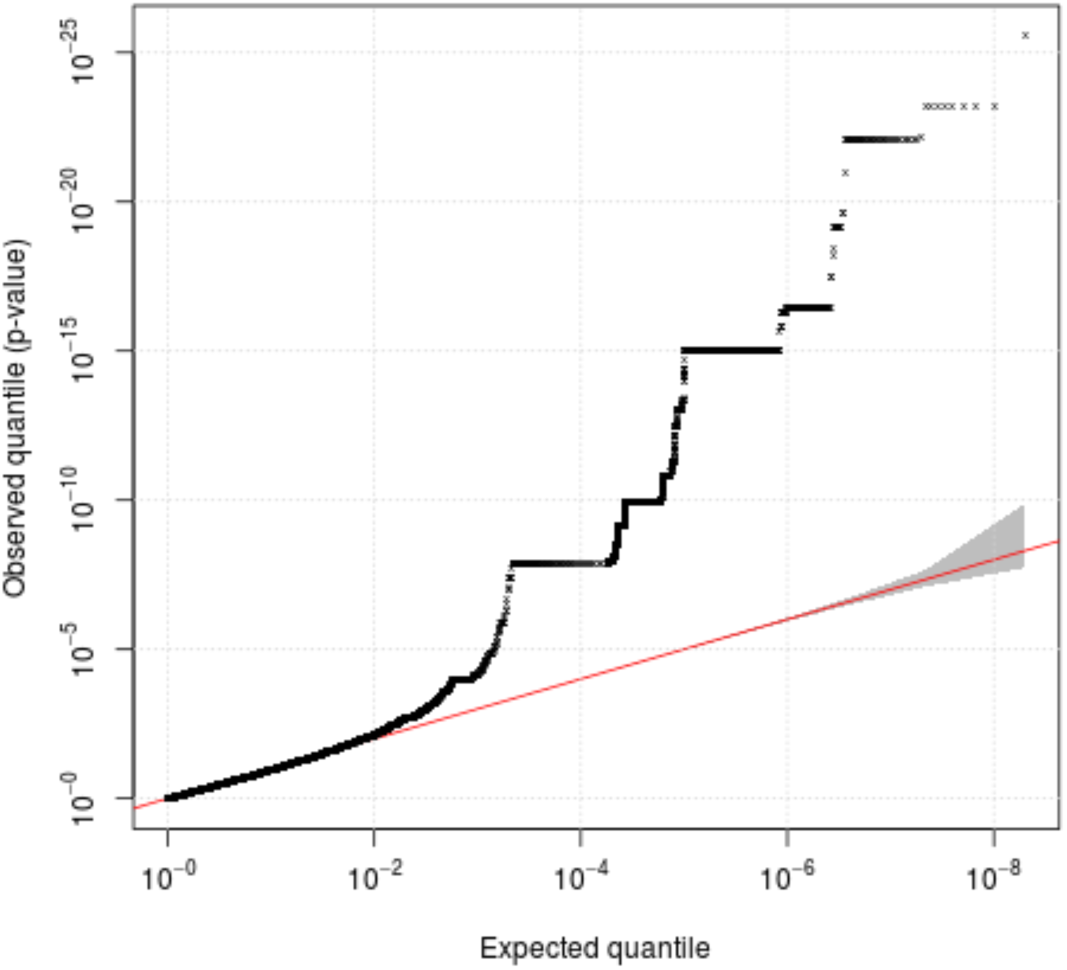
A) Distribution of Wald test P-values from 10^8^ simulations of genotypes for 77 cases and 2479 controls, assuming no true effect of genotypes on case/control status. P-values are plotted for both recessive and dominant inheritance models together (2x10^8^ P-values total). For each replicate, an allele frequency was drawn from a beta distribution with both shape parameters set to 0.8, roughly equivalent to the distribution seen on a SNP chip. Individual genotypes were then drawn from a binomial distribution based on this allele frequency, assuming Hardy-Weinberg equilibrium (HWE). Wald statistics were calculated as 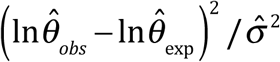 and evaluated against a chi-square distribution with 1 degree of freedom (equivalent to comparing the square root of this value to both tails of an N(0,1) distribution), where *θ*_*obs*_ is the observed OR from the appropriate 2x2 table (dominant or recessive inheritance model) after adding 0.5 each cell, and θ_exp_= [*e*_11_+0.5][*e*_12_+0.5]⁄[*e*_21_=0.5][*e*_22_=0.5], where {*e*_11_, *e*_12_, *e*_21_, *e*_22_} is the setoff expected genotype counts. Expected counts are produced using the number of cases and controls together with the minor allele frequency,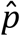, which is estimated from the full set of genotypes (e.g. for a recessive 2x2 table *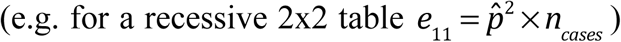*). The denominator of the Wald test statistic is 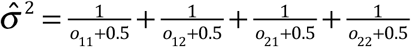 where {*o*_*11*_, *o*_*12*_, *o*_*21*_, *o 22*} is the observed set of genotype counts.

**Fig. S6.**
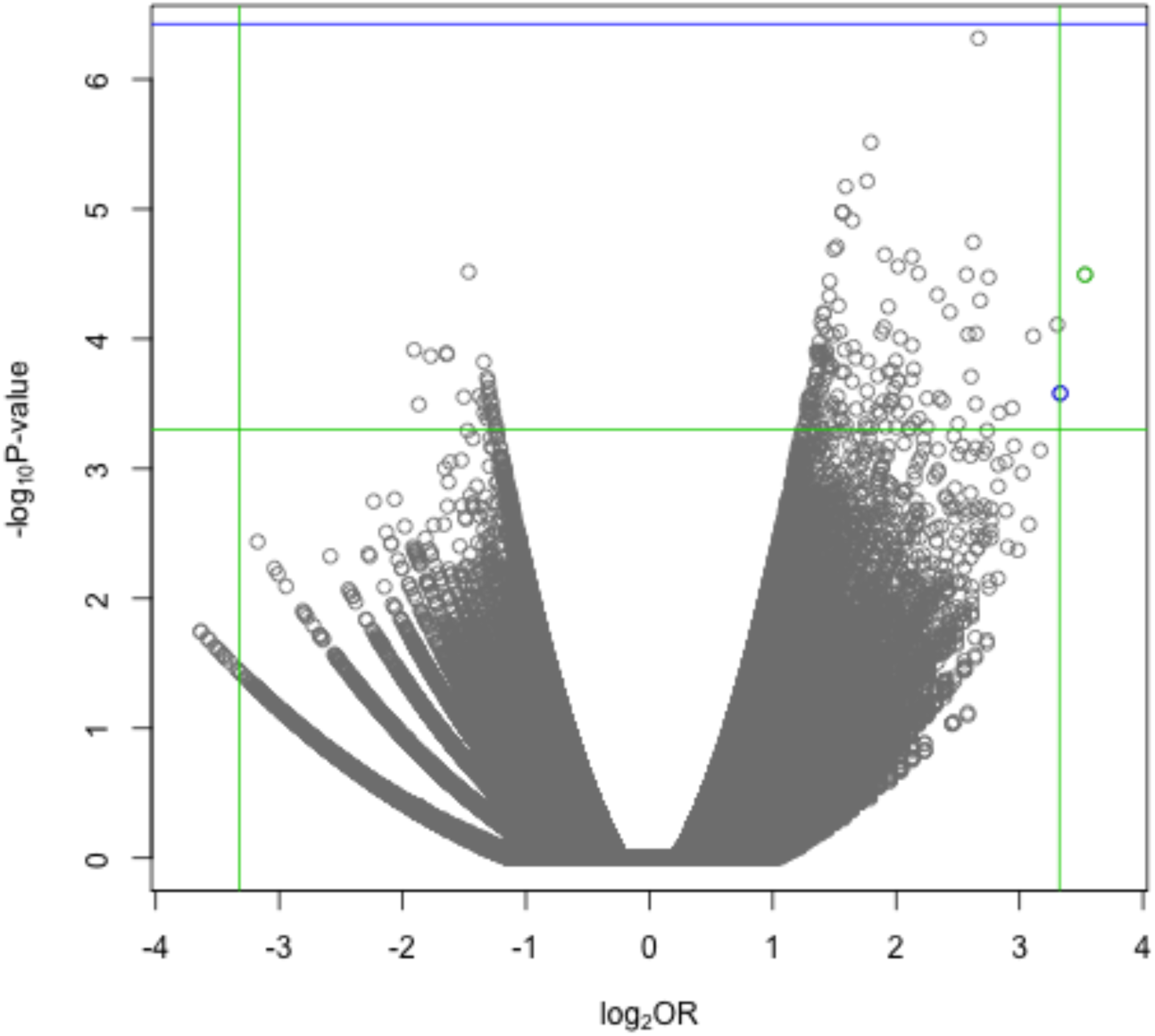
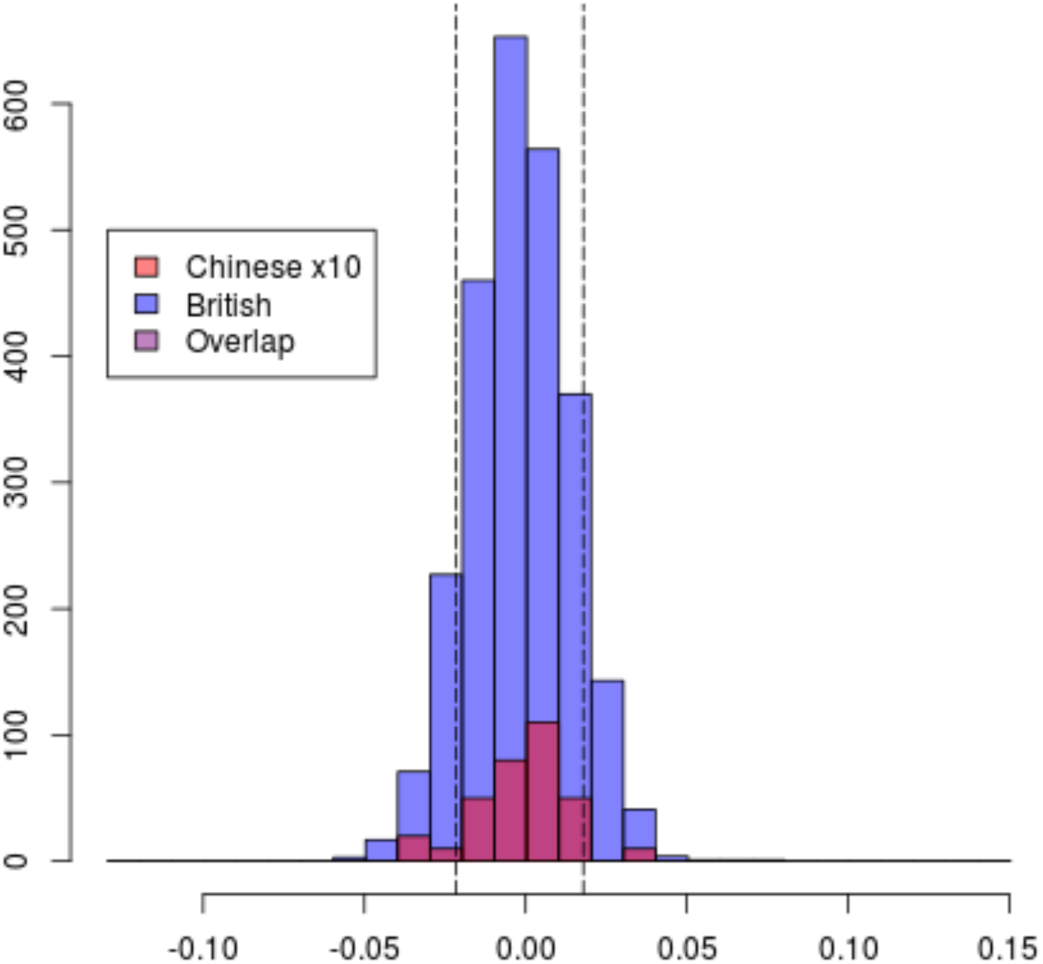
A)PC7, profile, females, upper extreme. The rs11642644 association is highlighted in green B) PC7, profile. Belonging to the upper 10% is associated with polymorphism at rs11642644 in females. The size of the Chinese histogram has been magnified by 10 times for visualisation purposes. Dotted lines show the upper and lower 10% quantiles

**Fig. S7.**
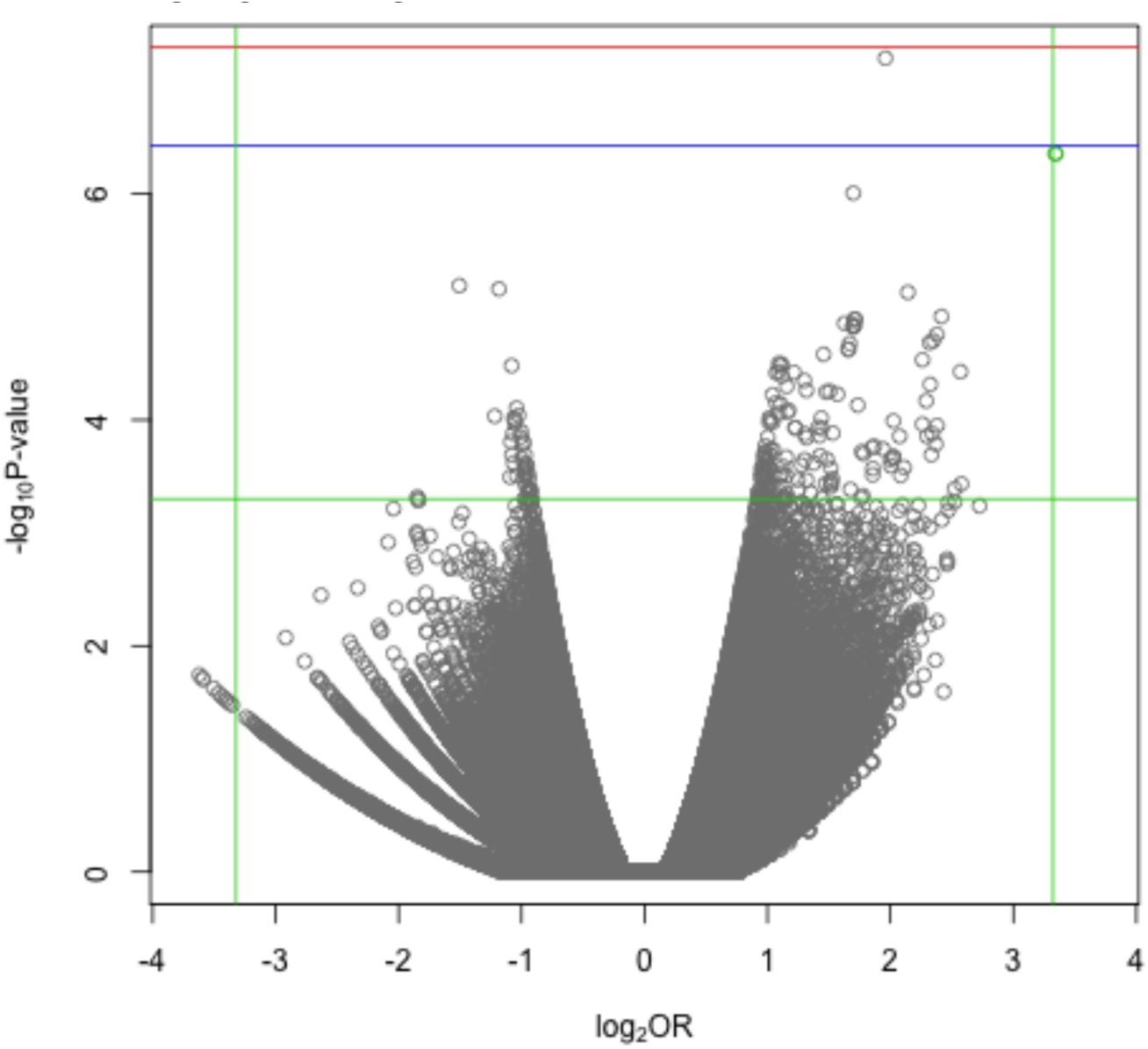
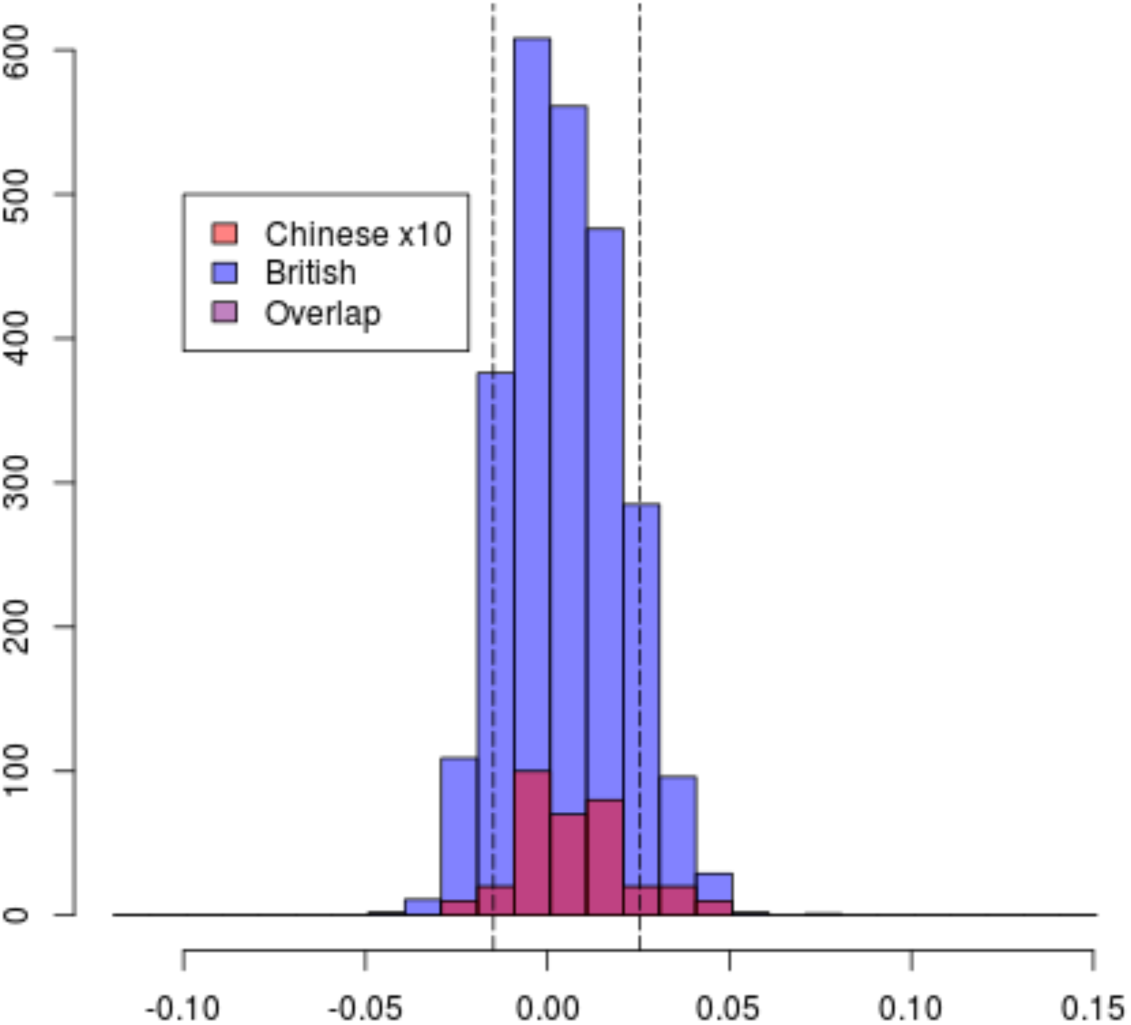
A) PC1, eyes, combined sexes, upper extreme. The rs7560738 association is highlighted in green. B) PC1, eyes. Belonging to the upper 10% is associated with polymorphism at rs7560738. The size of the Chinese histogram has been magnified by 10 times for visualisation purposes. Dotted lines show the upper and lower 10% quantiles

**Fig. S8.**
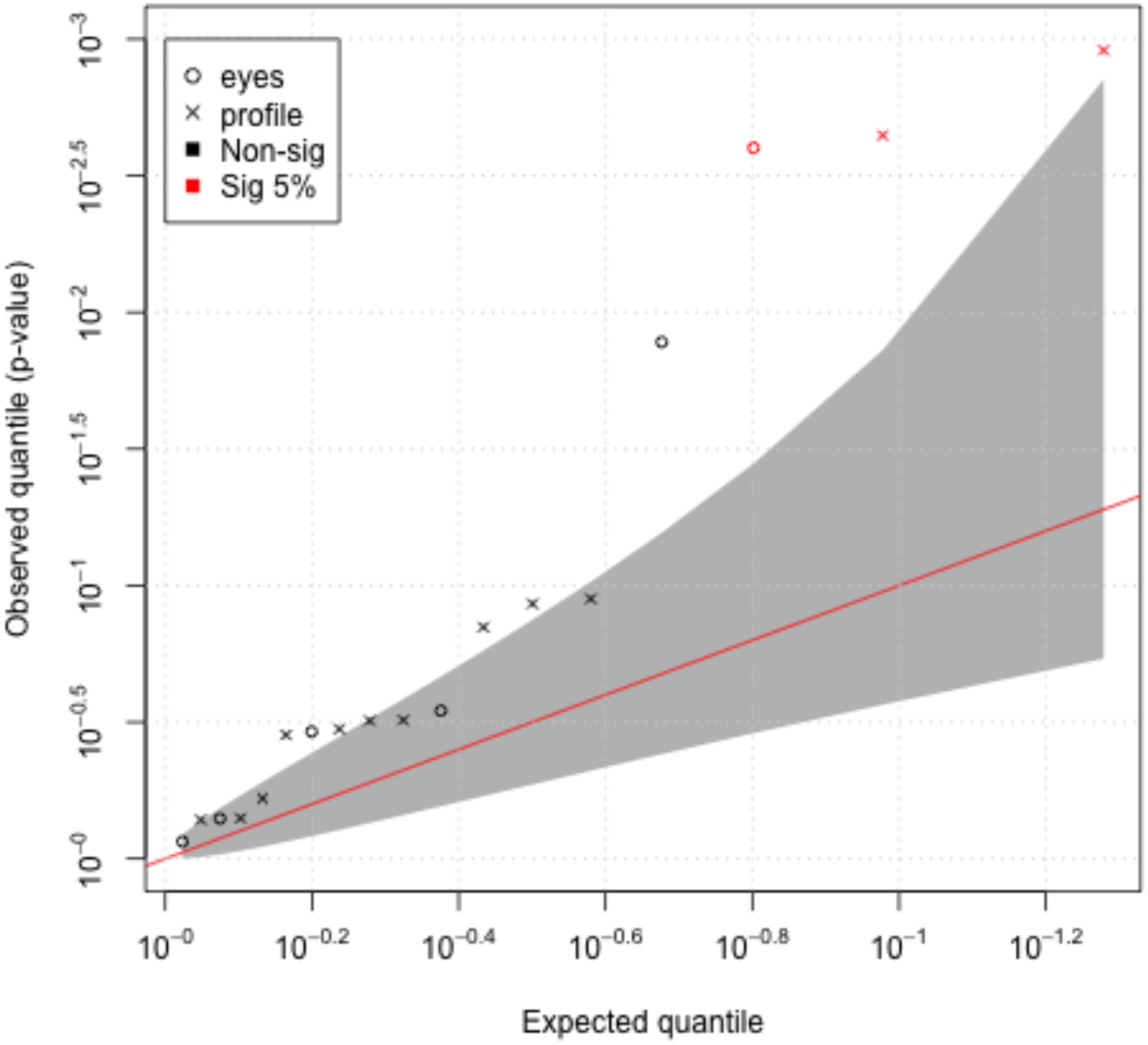
Quantile-quantile plot of the 18 replication P-values from permutation analysis of the TwinsUK data. The 95% confidence region is shown in grey, and SNPs passing an FDR of 5% are highlighted in red. Circles and crosses denote eyes and profile associations respectively.

**Fig. S9.**
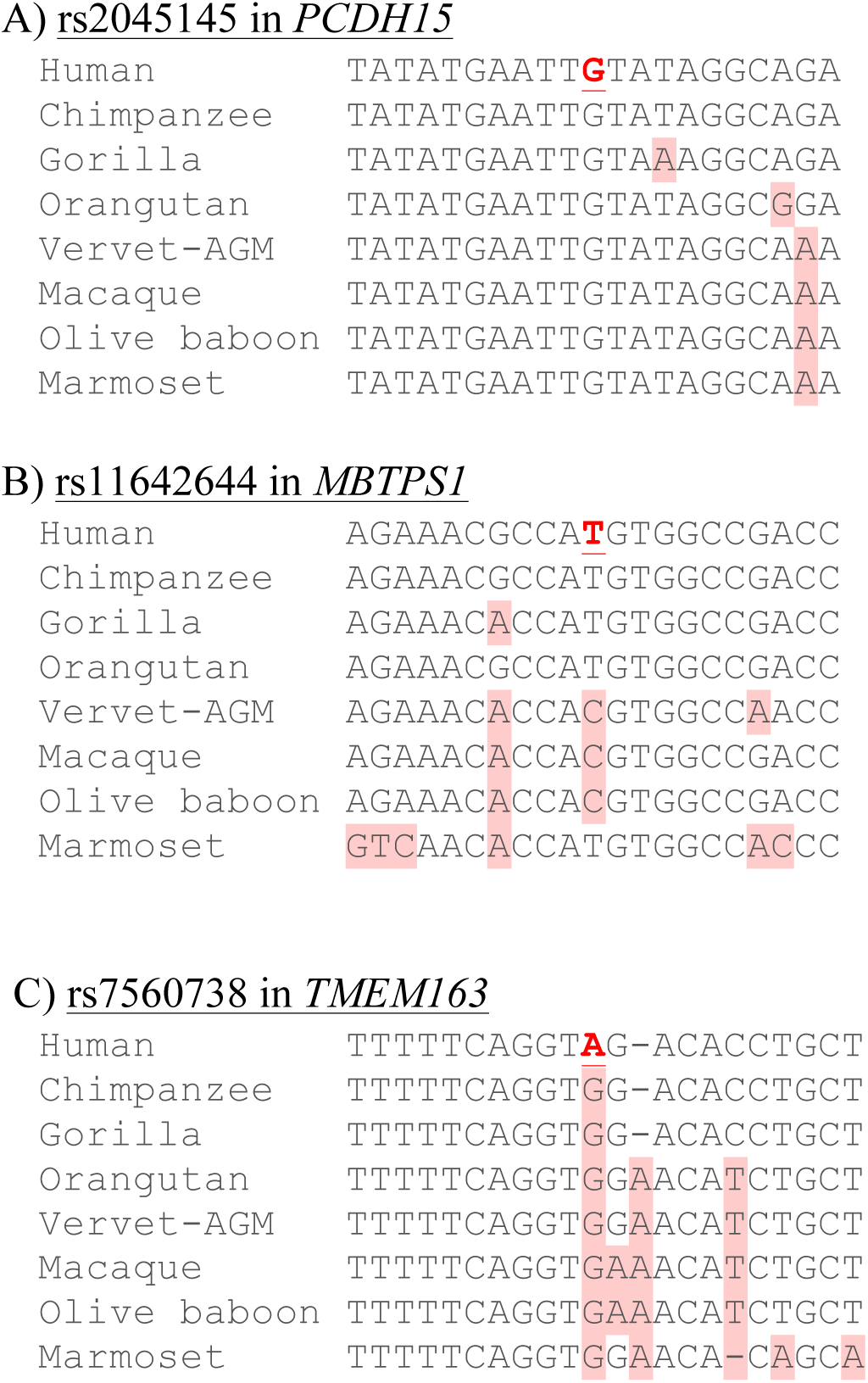
Conserved gene sequences around the discovery SNPs. In each case the base underlined in bold denotes the discovery SNP, and those on a red background differ from the human sequence.

